# A recombinant protein containing influenza viral conserved epitopes and superantigen induces broad-spectrum protection

**DOI:** 10.1101/2021.07.08.451596

**Authors:** Yansheng Li, Mingkai Xu, Yongqiang Li, Wu Gu, Gulinare Halimu, Yuqi Li, Zhichun Zhang, Libao Zhou, Hui Liao, Songyuan Yao, Huiwen Zhang, Chenggang Zhang

**Author notes:** Corresponding Author: Mingkai Xu. **Competing Interest Statement:** The authors declare that they have no known competing financial interests or personal relationships that could have appeared to influence the work reported in this paper.

## Abstract

Influenza pandemic poses public health threats annually for lacking vaccine which provides cross-protection against novel and emerging influenza viruses. Combining conserved antigens inducing cross-protective antibody response with epitopes activating cross-protective cytotoxic T-cells would offer an attractive strategy for developing universal vaccine. In this study, we constructed a recombinant protein NMHC consisting of influenza viral conserved epitopes and superantigen fragment. NMHC promoted the mature of bone marrow-derived dendritic cells and induced CD4^+^ T cells to differentiate into Th1, Th2 and Th17 subtypes. Mice vaccinated with NMHC produced high level of immunoglobulins which cross-bound to HA fragments from six influenza virus subtypes with high antibody titers. Anti-NMHC serum showed potent hemagglutinin inhibition effects to highly divergent group 1 (H1 subtypes) and group 2 (H3 subtype) influenza virus strains. And purified anti-NMHC antibodies could bind to multiple HAs with high affinities. NMHC vaccination effectively protected the mice from infection and lung damage challenged by two subtypes of H1N1 influenza virus. Moreover, NMHC vaccination elicited CD4^+^ and CD8^+^ T-cell responses to clear the virus from infected tissue and prevent virus spreading. In conclusion, this study provided proof of concept for triggering both B cells and T cells immune responses against multiple influenza virus infection, and NMHC may be a potential candidate of universal broad-spectrum vaccine for various influenza virus prevention and therapy.

## Introduction

Seasonal influenza viruses infected 5-15% of the population worldwide and killed several hundred thousand people every year in spite of the availability of antivirals and inactivated tetravalent vaccines, which are effective for most recipients(Ellebedy et al., 2014). As a cluster of RNA viruses, influenza viruses have segmented genome and error-prone RNA-dependent RNA polymerase which enable influenza viruses to undergo minor antigenic changes (antigenic drift) and major antigenic changes (antigenic shift)(Wang et al., 2010). This genomic mutability of influenza viruses permits the virus to evade adaptive immune response. The unpredictable variability of influenza A viruses causing yearly epidemics in human population, and time-consuming of current influenza vaccines industry for growth of the virus in chicken eggs are main reasons for lacking effective prevention against influenza infection up to date(Lu et al., 2014). Moreover, currently available vaccines induce antibodies mostly against homologous virus strains, but do not protect against antibody-escape variants of seasonal or novel influenza viruses. Therefore, development of a novel universal vaccine which could induce broad protection against various influenza viruses is urgent and important to overcome the problems caused by the annual epidemic of different types of influenza viruses. As to influenza A virus it is well recognized that conserved proteins or fragments of influenza A virus such as nucleoprotein (NP), matrix protein 2 ectodomain (M2e), hemagglutinin (HA2) stem (HA2)(Staneková and Varečková, 2010, El Bakkouri et al., 2011, De Filette et al., 2006), which could induce cross-protective immune response, are the most promising antigens to design a universal vaccine.

The hemagglutinin of influenza virus is a surface protein with strongly immunogenicity. Totally 18 HA subtypes of influenza A viruses could be classified into two groups (12 subtypes from group 1 such as H1 and H5, and 6 subtypes from group 2 such as H3 and H7) based on the phylogenetic relationships of HA proteins. HA is consisting of two subunits, membrane-distal globular domain HA1 and conserved stalk domain HA2 linked by a single disulfide bond(Wilson et al., 1981). The epitopes of broadly neutralizing antibodies (bnAbs) in the HA2 are more conserved across different influenza HA subtypes compared with the antigenic sites in the HA1(Julien et al., 2012, Ellebedy and Ahmed, 2012). The bnAbs induced by HA2 can against diverse influenza A virus subtypes by preventing fusion of the virus and host cell membranes (Julien et al., 2012). As a membrane protein, M2 protein is representing pH-gated proton channel incorporated into the viral lipid envelope, which is essential for efficient release of viral genome during virus entry(Schnell and Chou, 2008). The extracellular N-terminal domain of M2 protein (M2e), a 23 amino acid peptide, is highly conservative in influenza A strains and regarded as a valid and versatile vaccine candidate for inducing heterosubtypic antibody response against various human influenza strains. Nucleoprotein is a conserved inner antigen of influenza virus. Reports showed that NP is important to induce cellular immune response after natural infection. Furthermore, NP-specific helper T cells could augment protective antibody response and promote B cells to produce HA-specific antibodies(Townsend and Skehel, 1984). In brief, an effective vaccine against seasonal or pandemic influenza disease should contain both, conserved B-cell epitopes which act to prime the humoral immune system for a response capable of significantly diminishing virus replication immediately after infection, and T cell epitopes for the involvement of the cellular immune response to the overall protective effect(Moltedo et al., 2009, Hermesh et al., 2010).

Bacterial superantigen Staphylococcal Enterotoxin C2 (SEC2) has a notable ability to directly activate T lymphocytes without the presence of antigen-presenting cells (APC) and stimulate BMDCs maturation(Yao et al., 2018b, Dinges et al., 2000). Thus, on the one hand, SEC2 can activate T cells and BMDCs to produce large numbers of various cytokines such as interleukin-4 (IL-4) and interferon gamma (IFN-γ). On the other hand, SEC2 in extremely low dosage can induce CD8^+^ cytotoxic T lymphocyte (CTL) to specifically kill target cells such as virus-infected cells and tumor cells(Fu et al., 2020, Laidlaw et al., 2013). Furthermore, our previous study documented that SEC2 could link innate immunity and adaptive immunity through activating TLRs downstream signaling molecules including MyD88 and NF-κB, and could be served as a promising adjuvant for rabies vaccines to provide efficient protection against the lethal rabies virus exposure. Taking all these together, we hypothesize that SEC2 should have adjuvant or adjuvant-like effects to improve the protective efficiencies of influenza vaccine(Yao et al., 2018a).

In this study, to design a universal influenza vaccine, we connected the conserved fragments of NP (two epitopes of CTL and helper T lymphocytes), M2e, and highly conserved long α-helix regions of HA2 from group 1 (H1) and group 2 (H3 and H7), using flexible linker (GSAGSAG) to construct a fusion protein NMH as a subunit vaccine (Fig. 1A). To enhance the antigenicity of NMH, we fused SEC2 into the C-terminal of NMH to construct a conjugate vaccine NMHC. We evaluated influenza virus-specific antibodies-inducing ability of NMHC using direct-binding ELISA assay. Microneutralization-hemagglutination inhibition and cytotoxic lysis assay were performed to evaluate the bnAbs and immunogen induced by NMHC. And we also evaluated the universal protective efficiency of NMHC through protecting mice against influenza viruses in vivo.

**Figure 1.**
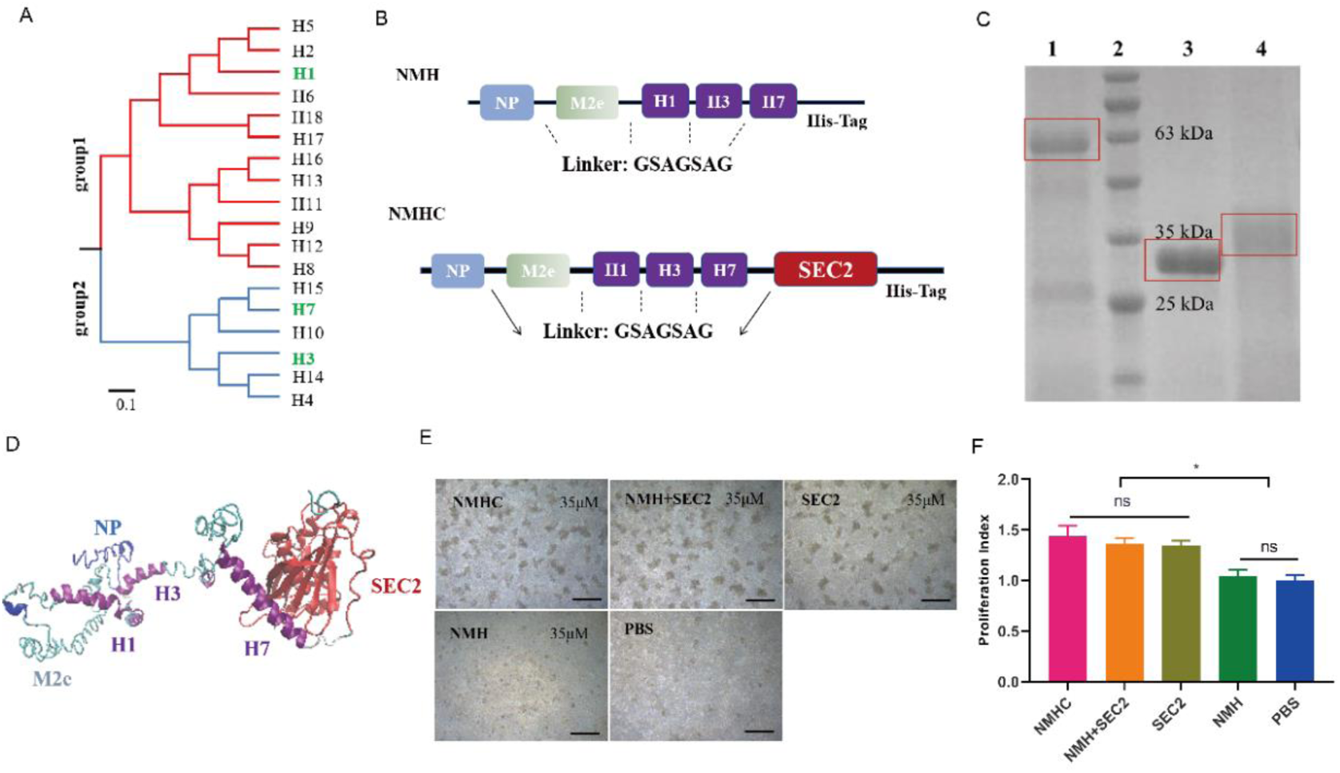
Design, purification, renaturation and verification of the recombinant protein NMHC. A, Phylogenetic tree of the 18 hemagglutinin subtypes of influenza A viruses based on amino acid sequences. Group 1 and group 2 subtypes are listed in red and blue respectively, and the HAs used in recombinant protein NMHC are listed in green. The amino acid distance scale bar denotes a distance of 0.1. B, NMH consists of the conserved gene segments of NP, M2e and HA2, which are connected by linker (GSAGSAG). NMHC design consists of NMH and linker followed by SEC2. C, NMHC, SEC2 and NMH purified and renatured from BL21(DE3) lysate. lane 1: renatured NMHC; lane 2: marker; lane 3: purified SEC2; lane 4: renatured NMH. D, the structure of the recombinant protein NMHC about homology modeling. The light blue, light green and purple portions are NP, M2e, HA2(H1, H3, H7) fragments respectively, and red portions fragment is SEC2 domain. E, murine splenocytes proliferation induced by NMHC, NMH+SEC2, SEC2, NMH and PBS were observed at 72 h. F, proliferation index was quantified by MTS assay. Scale bar = 200 μm. Data are represented as mean ± SD (n = 3). *, *p* < 0.05, ns, not significant. Source files of the gel used for the qualitative analyses was available in the Figure 1—source data 1.

## Results

### Construction, production and renaturation of recombinant proteins

The recombinant protein NMHC consisted of NMH at the N-terminal and SEC2 at the C-terminal, fused with a flexible linker sequence. To facilitate purification, a carboxyl-terminal His-tag reading frame was retained on the backbone of the expression vector pET-28a (+) (Fig. 1B). Recombinant proteins were expressed as inclusion body in plasmid-harboring *E. coli* strains and purified as a major band at 59 KDa (NMHC) and 31 KDa (NMH) in Coomassie blue-stained SDS-PAGE (Fig. 1C). After renaturation by dialysis, soluble recombinant proteins NMHC and NMH were got with purity >95% as confirmed by SDS-PAGE. The structural modeling of NMHC in silico revealed that the fused NMH was independent from the SEC2 region (Fig. 1D), which implied that the domains of NMH and SEC2 would not affect each other. To verify this hypothesis, splenocytes proliferation assay was performed. As shown in Fig. 1E, NMH did not induce any proliferation compared with PBS negative control, while SEC2, NMHC, and NMH+SEC2 induced obvious cell proliferative clusters in murine splenocytes at 72 h. The MTS result demonstrated that the PI of NMHC group was significantly higher than those of NMH and PBS (*p* < 0.05 Fig. 1F), albeit no significant difference was noted in these terms between the SEC2, NMH+SEC2, and NMHC groups.

### The biological activity of recombinant proteins in vitro

BMDCs are potent antigen-presenting cells (APCs) connecting the innate and adaptive immune responses. Maturation of murine BMDCs were evaluated by detecting the surface makers including CD80, CD86 and MHC II. Then, BMDCs (10^4^) were co-cultured with CD4^+^ T lymphocytes (10^5^) to evaluate the CD4^+^ T cells differentiation by flow cytometry. As showed in Fig. 2A, the expressions of CD80, CD86, and MHC II in NMHC, NMH+SEC2, SEC2 and NMH groups were significantly increased compared with PBS control group, which implied that the BMDCs maturation were induced by these treatments. BMDCs treated with NMHC significantly increased the expression of all these three molecules compared with NMH+SEC2 and SEC2 groups (*p* < 0.05). Furthermore, the expressions of CD80 and CD86 in NMHC and NMH+SEC2 group were significantly higher than that in NMH group (*p* < 0.01), albeit no statistical difference was noted between SEC2 and NMH+SEC2 groups (*p* > 0.05). As showed in Fig. 2B, the matured BMDCs induced by NMHC, SEC2, NMH+SEC2 and NMH significantly promoted CD4^+^ T cells to express Th1 cytokine IFN-γ and Th2 cytokine IL-4 compared with PBS control group (*p* < 0.01), and NMHC group exhibited higher expressions of IFN-γ, IL-4, and Th17 cytokine IL-17 than those in SEC2, NMH+SEC2 and NMH groups (*p* < 0.05 or *p* < 0.01). Moreover, no statistical difference was noted in any groups of regulatory T cell cytokine IL-10 expression (*p* > 0.05). This result implied that the mature BMDCs induced by NMHC could promote CD4^+^ T cell to differentiate or translated into Th1, Th2, Th17 cell subtypes but not Treg cells.

**Figure 2.**
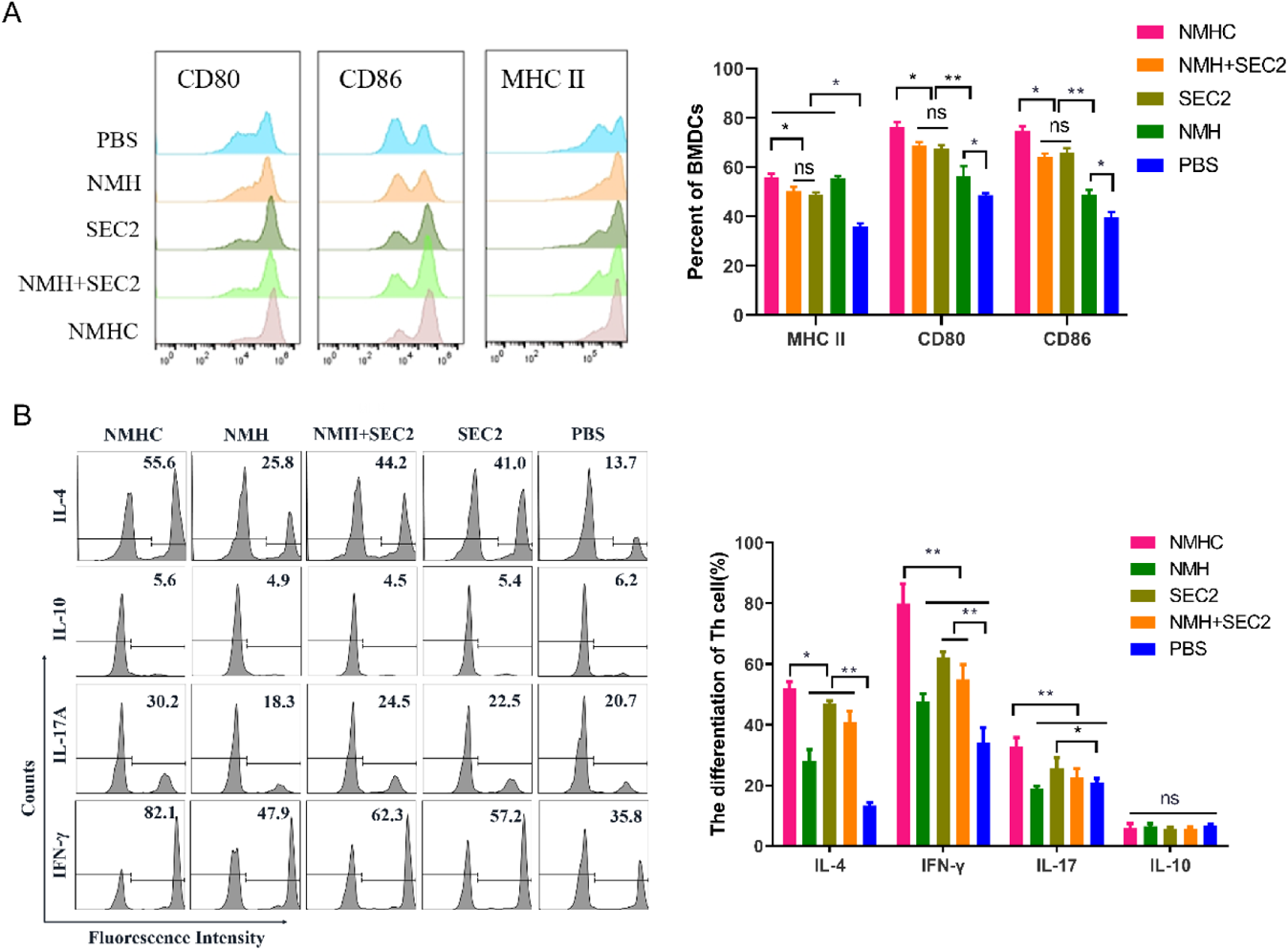
The expressions of co-stimulatory molecules CD80, CD86, and MHC II induced by recombinant proteins on BMDCs which inducted CD4^+^ T cell differentiation in vitro. A, BMDCs were treated with NMHC, NMH+SEC2, SEC2, NMH, and PBS as control for 48 h. Then BMDCs were stained with antibodies to MHC II, CD80, CD86 and analyzed with flow cytometry. B, Recombinant proteins-treated BMDCs (10^4^) were co-cultured with CD4^+^ T cells (10^5^) for 96 h, and CD4^+^ T cells differentiations were analyzed with flow cytometry. The results were showed with gated percentage. Data are represented as mean ± SD (n = 3). *, *p* < 0.05, **, *p* < 0.01, ns, not significant.

### Murine serum immunoglobulin isotyping elicited by recombinant proteins

Female BALB/c mice were immunized in a prime-boost-boost schedule with 2 weeks intervals. Serum samples were collected on the day 14, day 28, day 42, and day 100, and seven isotypes of κ immunoglobulins (accounted for 95% of immunoglobulin subtype in mice (Pricop et al., 1994)) were detected by CBA assay. As showed in Fig. 3, during 100 days post the first immunization, the IgG2a κ have significant changes. There is no statistical change for IgG2a κ in the first 14 days. While at day 28, NMHC, NMH, and NMH+SEC2 groups exhibited significantly enhanced production of IgG2a κ compared with the SEC2 and PBS groups (*p* < 0.05 for NMHC and NMH, and *p* < 0.01 for NMH+SEC2), and NMH+SEC2 treatment group showed the highest level of IgG2a κ. The similar results were detected at day 42, but the highest level of IgG2a κ was in NMHC group. Moreover, for long term effects, the production of IgG2a κ were returning in all the groups at day 100, but NMHC group still exhibited significantly enhanced IgG2a κ production compared with the PBS control group. There was no difference of IgG2a κ between the SEC2 group and PBS group at all the time points, which suggested that the IgG2a κ immunoglobulins detected in this study were specifically induced by NMH antigen but not by SEC2. Furthermore, the production of IgG2b κ showed observably enhanced in four treatment groups compared with control group at day 100 (*p* < 0.01), and there was no significant difference between each of the four groups (*p* > 0.05), probably because of the non-specificity antibody production. These data demonstrated that the vaccination acted as a productive immunogen boosting IgG2a which indicated T cell-dependent antibody production and suggested an affinity matured anti-virus response (Wang et al., 2010).

**Figure 3.**
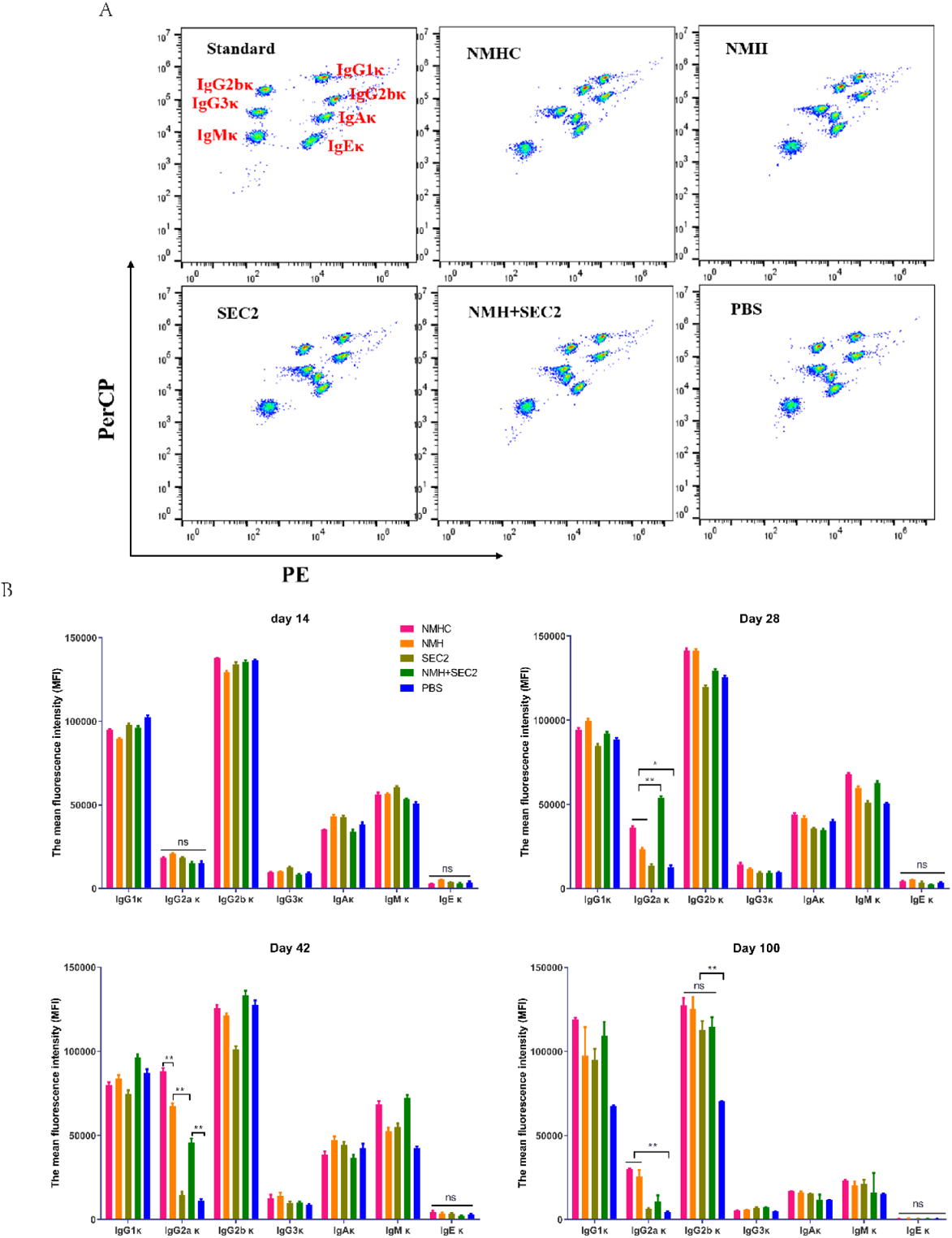
Murine serum immunoglobulin isotyping elicited by recombinant proteins. Sera were taken from immunized mice on the day 14, day 28, day 42, and day 100 after immunization, and immunoglobulins were examined by Cytometric Bead Array. A, seven clusters of beads represented the immunoglobulins of IgG1κ, IgG2a κ, IgG2b κ, IgG3 κ, IgA κ, IgM κ and IgE κ from top to bottom when detected in FL2 (PE) and FL3 channels (PerCP). B, the contents of immunoglobulins in different time points were represented by median fluorescence intensity, Data are represented as mean ± SD (n = 3). *, *p* < 0.05, **, *p* < 0.01, ns, not significant.

### Recombinant proteins induced neutralizing antibody in mice

Because the stem-directed bnAbs mediated neutralization by inhibiting membrane-fusion but not by blocking the virus from binding to receptors on the host cells(Julien et al., 2012), the standard hemagglutinin inhibition assay is not fit for this study (as in Table S1, neutralizing antibody was not detected in serum at the lowest dilution 1:8). To evaluate the titers of neutralizing antibody induced by recombinant proteins, an optimized Microneutralization-Hemagglutinin inhibition assay were performed as described in methods above. As showed in Fig. 4, NMHC, NMH+SEC2, and NMH induced high levels of in neutralizing antibody titers following the third immunization, which provide a critical role in curbing H1N1 A/Michigan/45/2015 (group 1) and H3N2 A/Hong Kong/ 4801/2014 (group 2) viruses replication, compared with SEC2 and PBS. Furthermore, NMHC and NMH+SEC2 exhibited higher titers than NMH (Fig. 4A and B). This result suggested that sera from recombinant proteins immunized mice had substantial heterosubtypic neutralizing activity, and the mechanism of protection might be due to inhibiting virus-host cell membrane fusion(Okuno et al., 1994).

**Figure 4.**
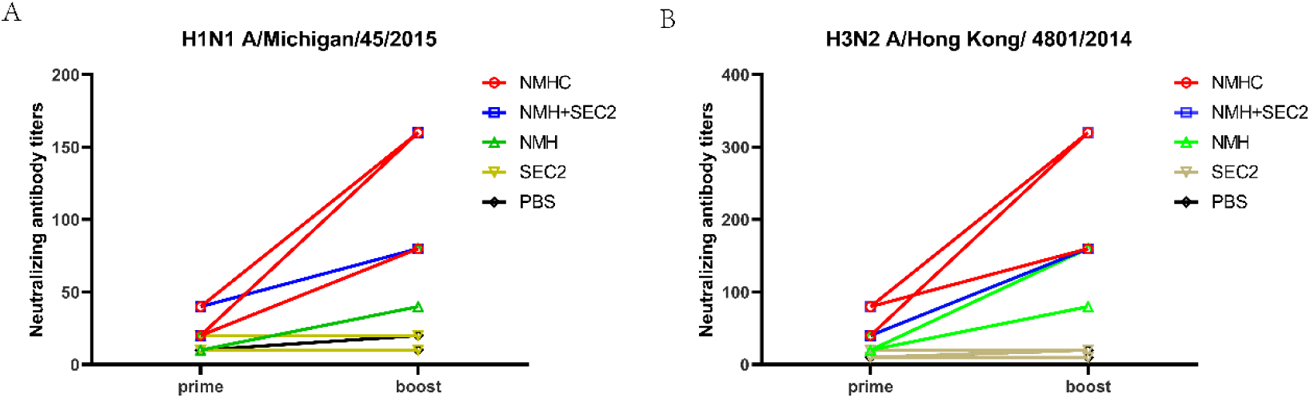
Recombinant proteins elicit broadly cross-reactive bnAbs in mice. Sera were taken from immunized mice on day 14 (prime) and day 42 (boost) after immunization, and the neutralization assays were performed against H1N1 (A) or H3N2 (B) influenza viruses. The titers of each serum sample were defined as the reciprocal of the highest dilution.

**Table 1.**
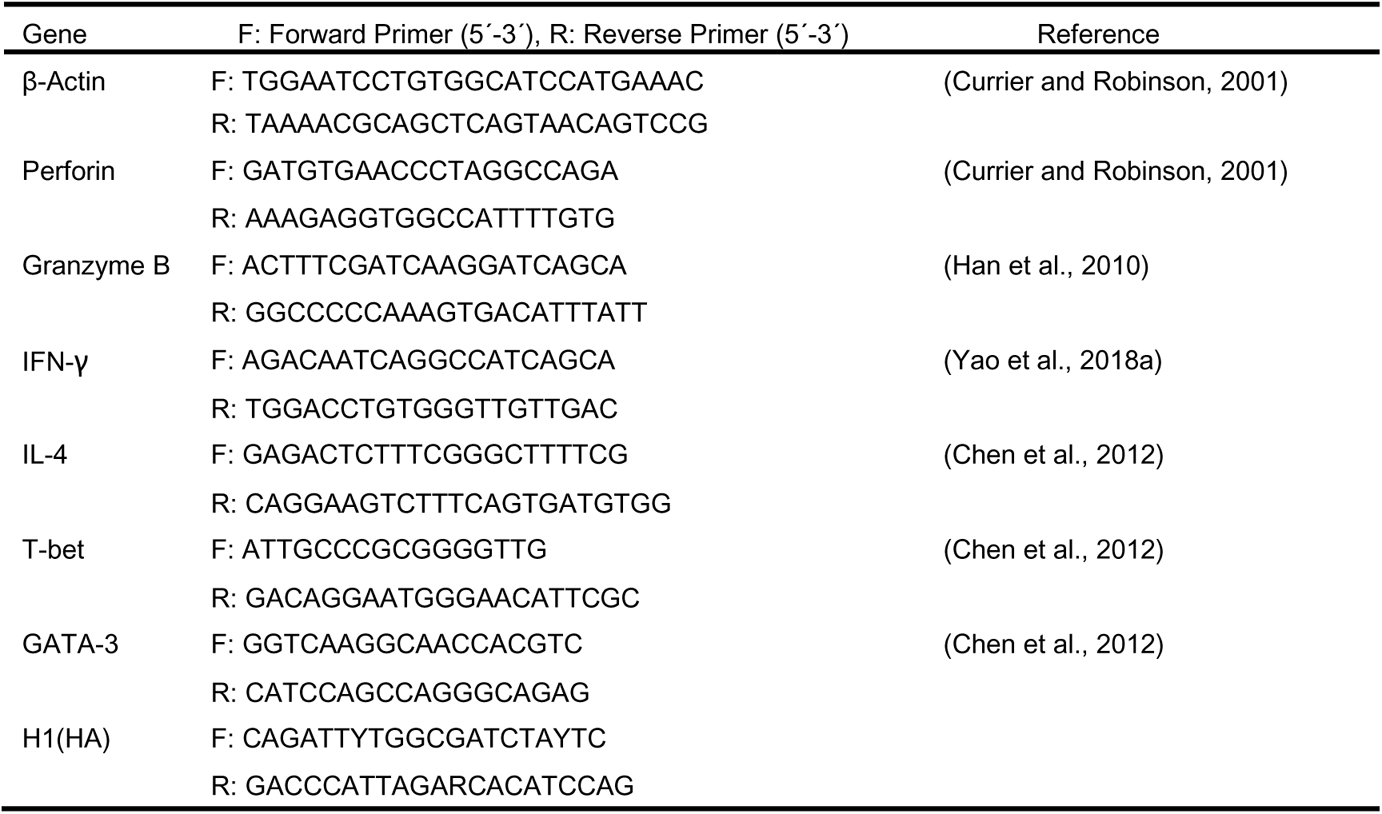
Sequences for qPCR primers

### Recombinant proteins induced binding antibodies to a broad panel of HA subtypes

To determine whether immunization with recombinant proteins could induce broadly binding antibodies, ELISA assays were performed with four recombinant HAs and two split virions. As expected, NMHC, NMH+SEC2, and NMH induced high levels of antibodies against various HA subtypes, while SEC2 and PBS did not induce any effective antibodies (Fig. 5A-F and Table S2). NMHC and NMH+SEC2 have more potencies than NMH to induce H3, H5, and H7 specific antibodies (Fig. 5C-E), and NMHC exhibited higher ability to induce H1 specific antibody than NMH+SEC2 and NMH (Fig. 5A). Moreover, taken the strains of MI/45(H1) and HK/4801(H3) for examples, we also measured influenza-specific IgG1 and IgG2a subclasses of antisera, which were essential to understand the B cell somatic hypermutation and subclass-switching(Quan et al., 2007). The results showed that NMHC induced higher H1N1-specific IgG1 and IgG2a than NMH and NMH+SEC2 (Fig. 5G and H), and NMHC and NMH+SEC2 induced higher H3N2-specific IgG1 and IgG2a than NMH (Fig. 5I and J), which were consistent with the results of total IgG as showed in Fig. 5A and Fig. 5C. These data indicated that the recombinant vaccine NMHC could significantly induce broad binding or neutralizing antibodies activity and might confer cross-subtype protection.

**Figure 5.**
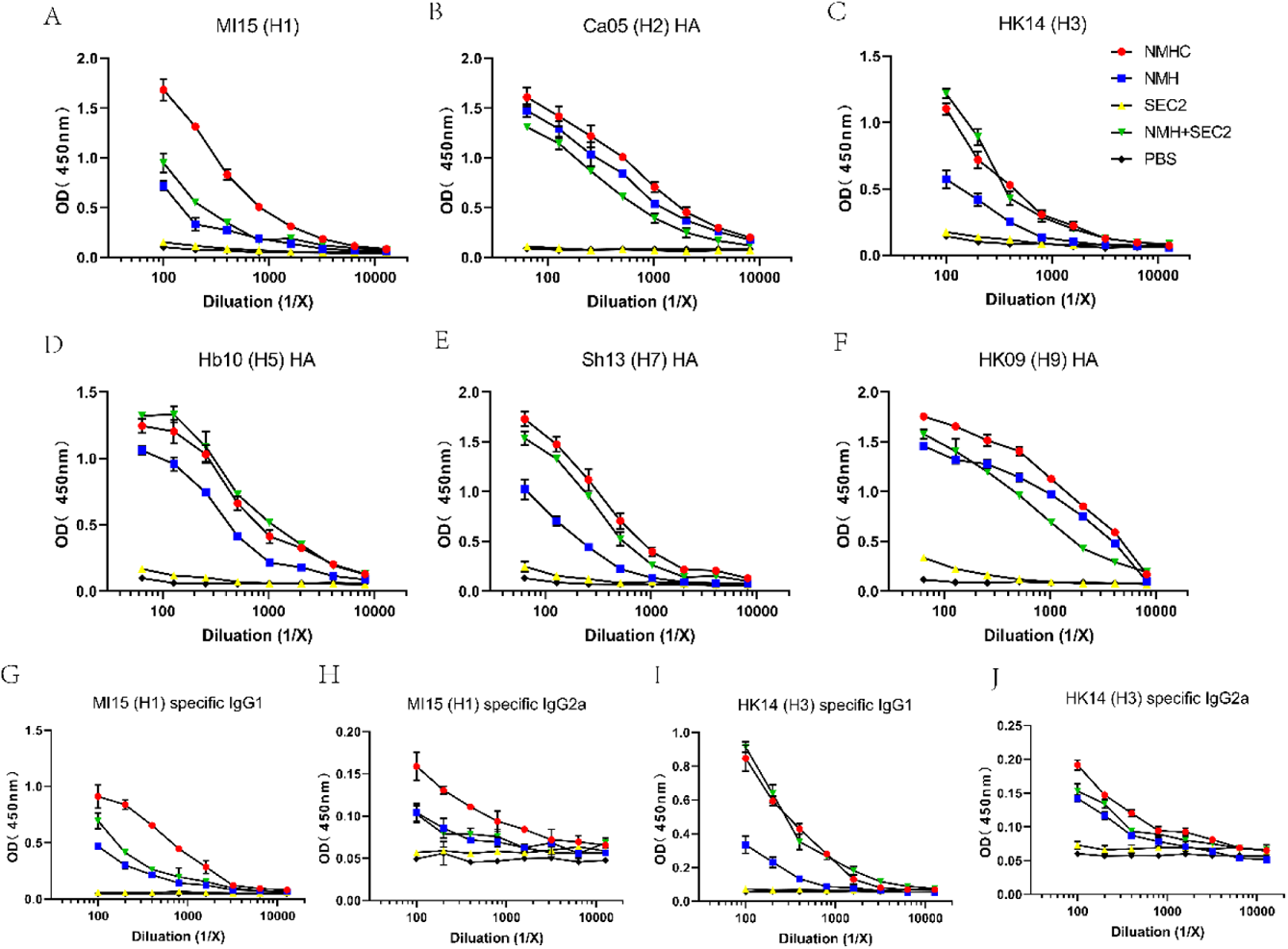
Breadth of the antibody response elicited by the recombinant protein was determined by ELISA of the pooled antisera against purified rHA proteins or split virion. A-F, total IgG against (A) H1N1 A/Michigan/45/2015, (B) H2N2 A/Canada/720/2005, (C) H3N2 A/Hong Kong/ 4801/2014, (D) H5N1 A/Hubei/1/2010, (E) H7N9 A/Shanghai/2/2013, (F) H9N7 A/Hong Kong/35820/2009. G-J, IgG1 or IgG2a specific against split virion of influenza viruses. (G) IgG1 specific against H1N1 A/Michigan/45/2015, (H) IgG2a specific against H1N1 A/Michigan/45/2015, (I) IgG1 specific against H3N2 A/Hong Kong/4801/2014, (J) IgG2a specific against H3N2 A/Hong Kong/ 4801/2014.

### NMHC induced antibodies with high binding affinity to various HAs

We purified the serum antibodies and evaluated the ability of serum antibodies to form a stable complex with the NMH antigen. The results of pull-down assay showed that the antibodies purified from NMHC, NMH+SEC2 and NMH antisera could bind and pull down NMH antigen, while SEC2 and PBS could not (Fig. S1). Subsequently, SPR experiments were performed to quantify the binding affinities of antibodies purified from anti-NMHC sera. NMHC-induced Abs displayed potent binding affinity to H1, H2, H3 and H5 fragments with KD value of 98 nM, 73.9 nM, 685 nM, and 72.9 nM, respectively (Fig. 6). This result implied that immunized with NMHC could elicit broadly cross-reactive protection against various influenza viruses.

**Figure 6.**
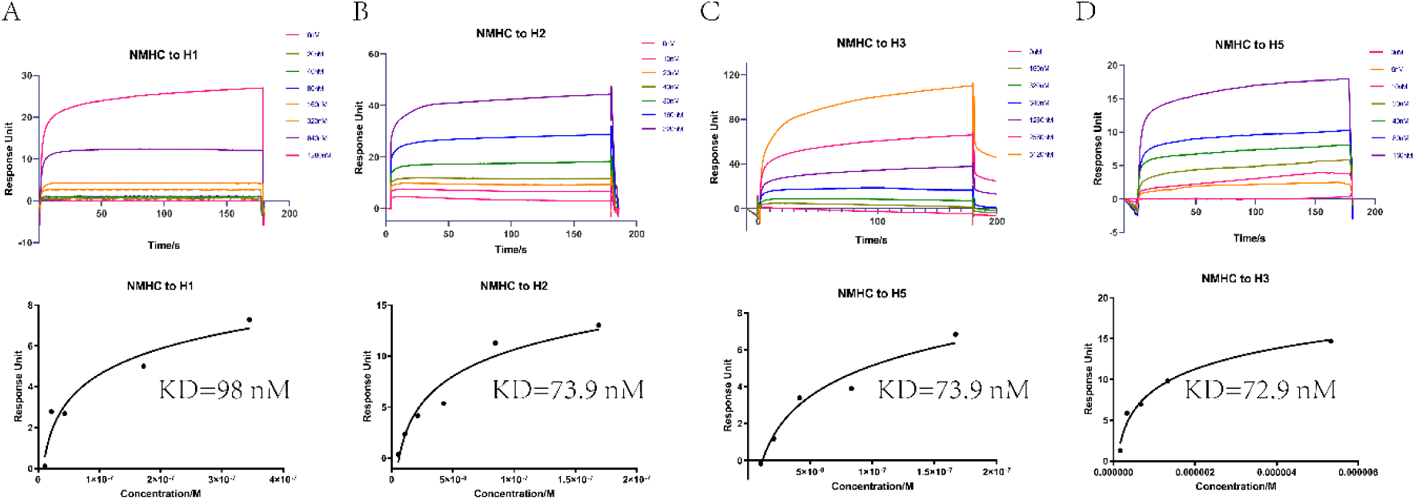
Affinities of HAs to antibodies purified from anti-NMHC sera. The overlays of binding kinetics of the HAs at different concentrations and the affinity of KD values, determined with Biacore were shown. (A) split virion of H1N1 A/Michigan/45/2015, (B) H2N2 A/Canada/720/2005, (C) split virion of H3N2 A/Hong Kong/ 4801/2014, (D) H5N1 A/Hubei/1/2010. The KD values were calculated using a steady affinity state model by the Biacore T200 evaluation software (Version 3.1). Source files of the overlays of binding kinetics used for the affinity analyses were available in the Figure 6—source data 2.

### Recombinant proteins induced ASC responses

ELISPOT assays were performed to measure recombinant proteins-induced anti-H1N1, H3N2, and NMH specific B-cell responses (Fig. 7A). It is important to note that H1N1 and H3N2 viruses are the components of seasonal vaccines for the 1978–2020 influenza seasons. Firstly, we determined the kinetics of anti-H1N1 and H3N2 B cell responses in splenocytes from pre- and post-immunized mice (Fig. 7B and C). The responses were positive correlation with vaccination times and peaked at day 42. At the peak of the response, the frequencies of NMH, H1N1, and H3N2 specific ASCs were significantly higher in NMHC, NMH+SEC2, and NMH immunized groups than those in SEC2 and PBS treated groups (*p* < 0.01, Fig. 7D, E and F), and there was no statistical difference between the SEC2 and PBS groups (*p* > 0.05). Moreover, at the peak point, the frequencies of NMH, H1N1, and H3N2 specific ASCs induced by NMHC were 73±5.2, 48±5.7, and 64±2.1 per million, respectively, which were about 1.5-fold higher than the frequencies of these three ASCs induced by NMH (*p* < 0.05, Fig. 7D, E and F). Notably, NHMC immunization induced higher frequencies of H1N1 and H3N2 specific ASCs than NMH+SEC2 immunization did (*p* < 0.05, Fig. 7E and F). Interestingly, in comparison to the prime vaccination, we observed a large increase in the frequency of anti-H1N1 and H3N2 ASCs at day 100 in NMHC immunized group (Fig 7B and C), which implied that NMHC probably induce cross-reactive memory B cells after three times of immunization(Purtha et al., 2011). In summary, NMHC could effectively induce B-cell responses and elicit serological memory that might be maintained by long-lived antibody-secreting plasma cells and reinforced by memory B cells, which could rapidly differentiate into antibody secreting cells when exposed with antigen again.

**Figure 7.**
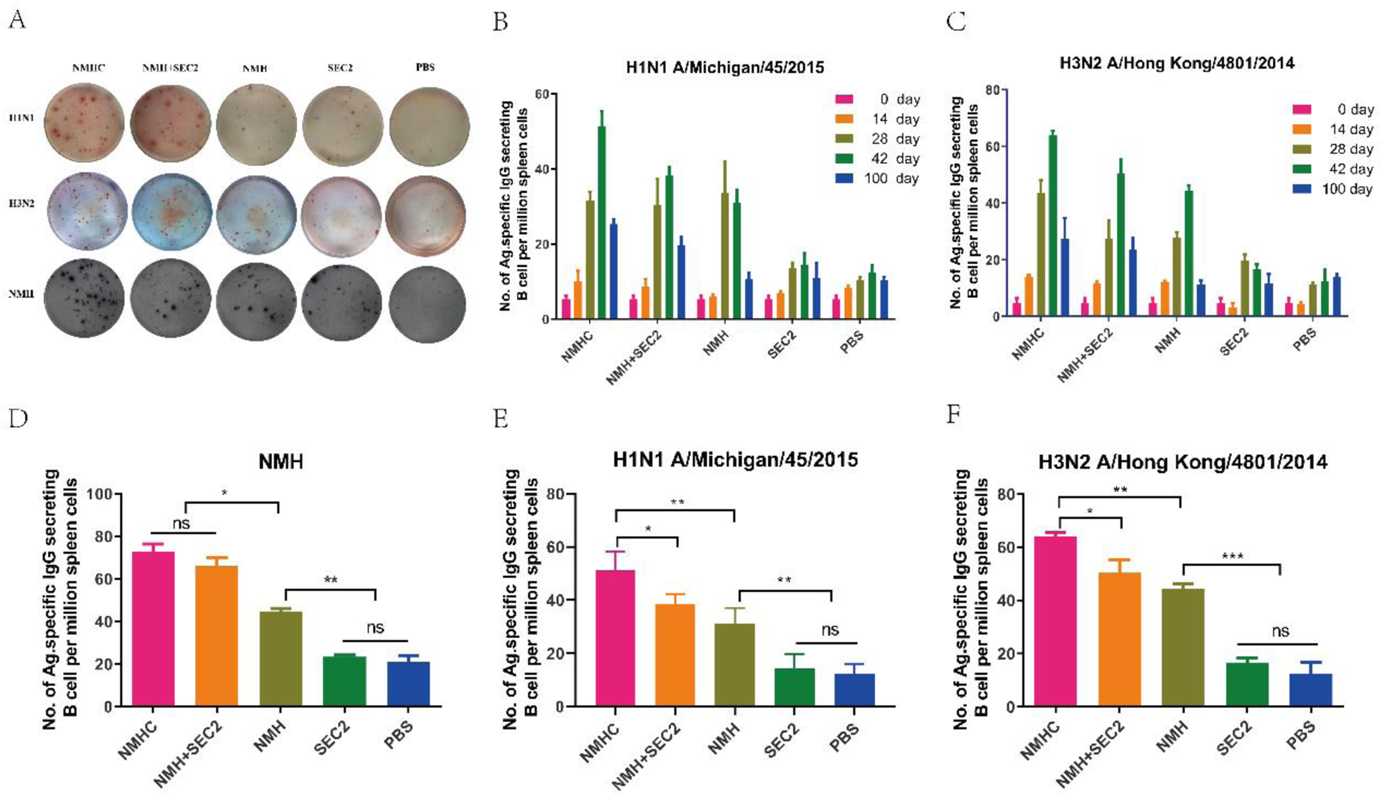
The specific antibody secreting cells response to H1N1, H3N2 and NMH were measured with ELISPOT assay. Splenocytes were isolated and collected from immunized mice on the day 14, day 28, day 42 and day 100. The cells were added in ELISPOT wells, which coated with the NHM proteins, H1N1 or H3N2 influenza viruses. A, each spot in the well represented an ASC. B and C, the frequency of H1N1-specific and H3N2-specific antibody secreting cells on day 0, day 14, day 28, day 42 and day 100. D, E and F, specific antibody secreting cells response to NMH, H1N1 and H3N2 influenza viruses on day 42. Data are represented as mean ± SD (n = 3). *, *p* < 0.05, **, *p* < 0.01, ***, *p* < 0.001, ns, not significant.

### Recombinant proteins induced T Cell responses

We studied the vaccine-induced CD4^+^ and CD8^+^ T cells immune responses which were potent mediators of heterosubtypic immunity against different influenza viruses. Since IFN-γ and IL-4 were typically produced by Th1 and Th2 cells, respectively(Yao et al., 2018a), ELISPOT assays were performed to measure IFN-γ or IL-4-secreing cells induced by recombinant proteins immunization. The results showed that immunized with NMHC and NMH+SEC2 significantly induced the upregulation of IFN-γ or IL-4-producing splenocytes (*p* < 0.01 to PBS group; Fig. 8A), and there was no statistical difference compared with SEC2 group (*p* > 0.05). Furthermore, immunized with NMH alone did not induce any cytokines-producing splenocytes compared with PBS group (*p* > 0.05). Additional, IFN-γ and IL-4 as well as the specific transcription factors T-bet and GATA3 derived from Th1 and Th2 respectively were quantified by qPCR. The results were consistent with those from ELISPOT assay that NMHC, NMH+SEC2, and SEC2 significantly induced the upregulation of IFN-γ and IL-4 transcription compare with NMH alone and PBS (*p* < 0.05), and the similar changes were found in T-bet and GATA3 transcription factors. Taken together, NMHC immunization could activate both Th1 and Th2 cells which were necessary for immune responses to clear virus infection.

**Figure 8.**
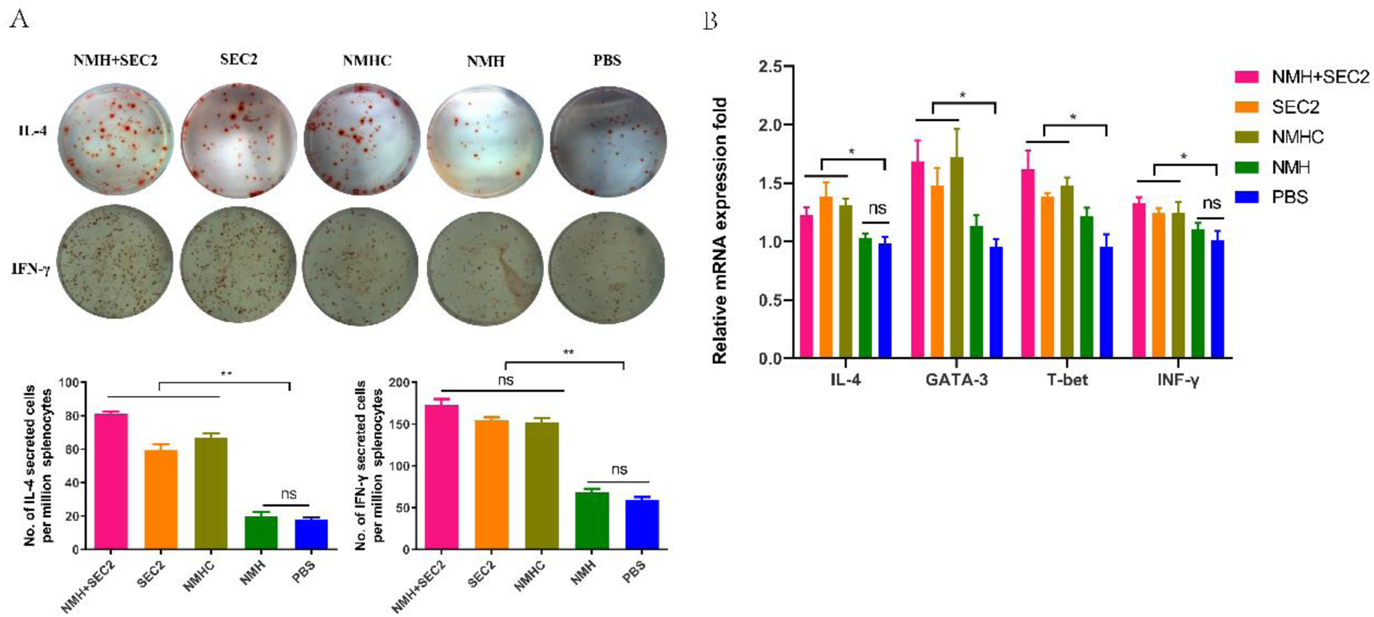
Recombinant proteins induced CD4^+^ T cell responses. The IFN-γ and IL-4 cytokines producing cells were measured with ELISPOT assay. A, splenocytes were isolated from immunized mice in each group on days 42, and the cells were plated in ELISPOT wells, which were coated with anti-IFN-γ or anti-IL-4 antibodies. Secreted cells responded to vaccine stimulation were measured with ELISPOT assay. B, the mRNA levels of IFN-γ, IL-4, T-bet, and GATA3 were measured by qPCR. Results are expressed as the mean value ± SD (n=3) *, *p* < 0.05, **, *p* < 0.01, ns, not significant.

The cytotoxicity effects of CD8^+^ T cells are important to clear virus infected cells, which play the key roles for the efficient control of influenza virus spreading(Yao et al., 2018a). To assess if the recombinant vaccines could augment antigen-specific CD8^+^ T cell responses, the equal number of CFSE ^high^ target cells and CFSE ^low^ non-target control cells were mixed and injected intravenously into recipient mice for twelve hours and the specific CTL were examined in vivo. Firstly, we noted that Alexa Fluor 488-labeled NMH antigens could be effectively internalized into splenocytes isolated from naïve mice detected by LSCM (Fig. 9A). Secondly, target cells and control cells were easily distinguished when labeled with 10 μM or 1 μM of CFSE, and the fluorescence intensity of the former was higher than the latter (Fig. 9B). The result showed that the relative number of residual target cells (CFSE ^high^) isolated from recipient mice immunized with NMHC and NMH+SEC2 were drastically reduced compared with NMH, SEC2 and PBS treatment (*p* < 0.01; Fig. 9C). Furthermore, NMHC significantly induced perforin and granzyme express in transcription level, which also confirmed that the NMHC could induce strong antigen-specific cytotoxic responses (Fig. 9D).

**Figure 9.**
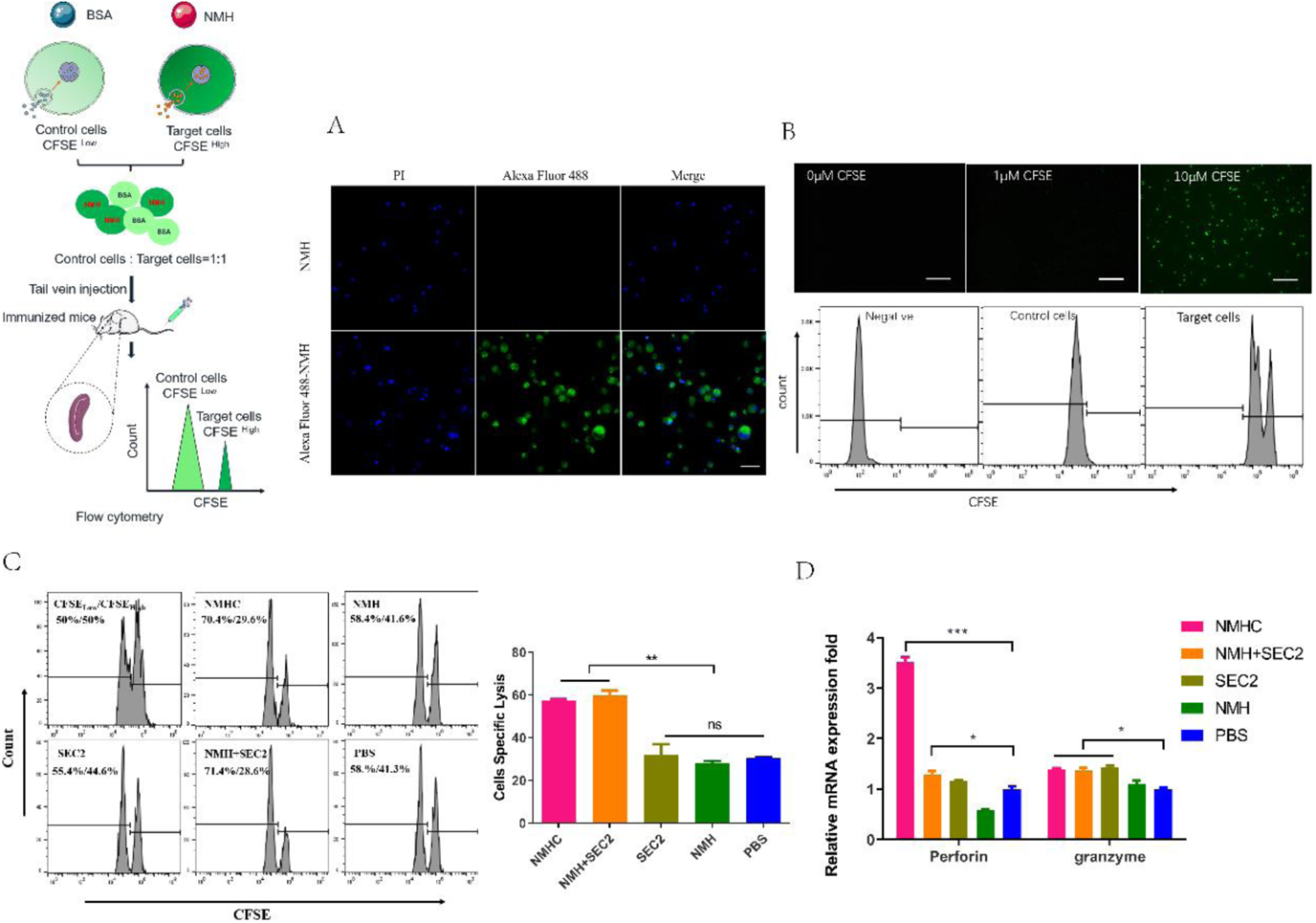
Effects of the recombinant protein NMHC on CD8^+^ T cell responses. A, to verify that NMH antigens can be effectively internalized into splenocytes, splenocytes prepared from naïve BALB/c mice were co-cultured with Alexa Fluor 488-labeled NMH or NMH alone and detected by fluorescent inverted microscope. B, then, fresh isolated naïve splenocytes were divided into two parts. One part was pulsed with 5 μg/mL NMH, labeled with 10 μM of CFSE and termed as CFSE high target cells, the other part was loaded with BSA, labeled with 1 μM of CFSE and defined as CFSE low control cells, both of them detected by fluorescent inverted microscope and flow cytometry. C, the equal number of two parts cells were mixed and injected into recipient mice. Twelve hours later, the splenocytes from these mice were used to examine the antigen specific cytolytic responses. D, the mRNA levels of perforin and granzyme of recipient mice were examined by qPCR. Statistical analyses were performed using Student’s t-test and one-way ANOVA. *, *p* < 0.05, **, *p* < 0.01, ***, *p* < 0.001, ns, not significant.

### Immunized with recombinant proteins provide in vivo protection against influenza virus challenge

To evaluate whether the recombinant protein immunization conferred cross-strain protection, mice were challenged with virus strains of Bris/02(H1) or MI/45(H1) at two weeks after the third immunization. At 6 days post-infection, the qPCR analysis of the lung tissues demonstrated that the mice immunized with NMHC exhibited up to 4 logs reduction in viral NP gene copy number of Bris/02(H1) and MI/45(H1), compared with the PBS treated groups (Fig. 10A and B). And mice immunized with NMH+SEC2 got the similar results. During the whole challenge experiment, only PBS treated groups showed reversible loss of body weights (Fig. 10C and D). The histopathological examination demonstrated that there were no visible pathological changes in the lungs from the NMHC-immunized mice at 6 days post-infection, while the PBS control group exhibited severe alveolar damage and interstitial inflammatory infiltration (Fig. 10E and F).

**Figure 10.**
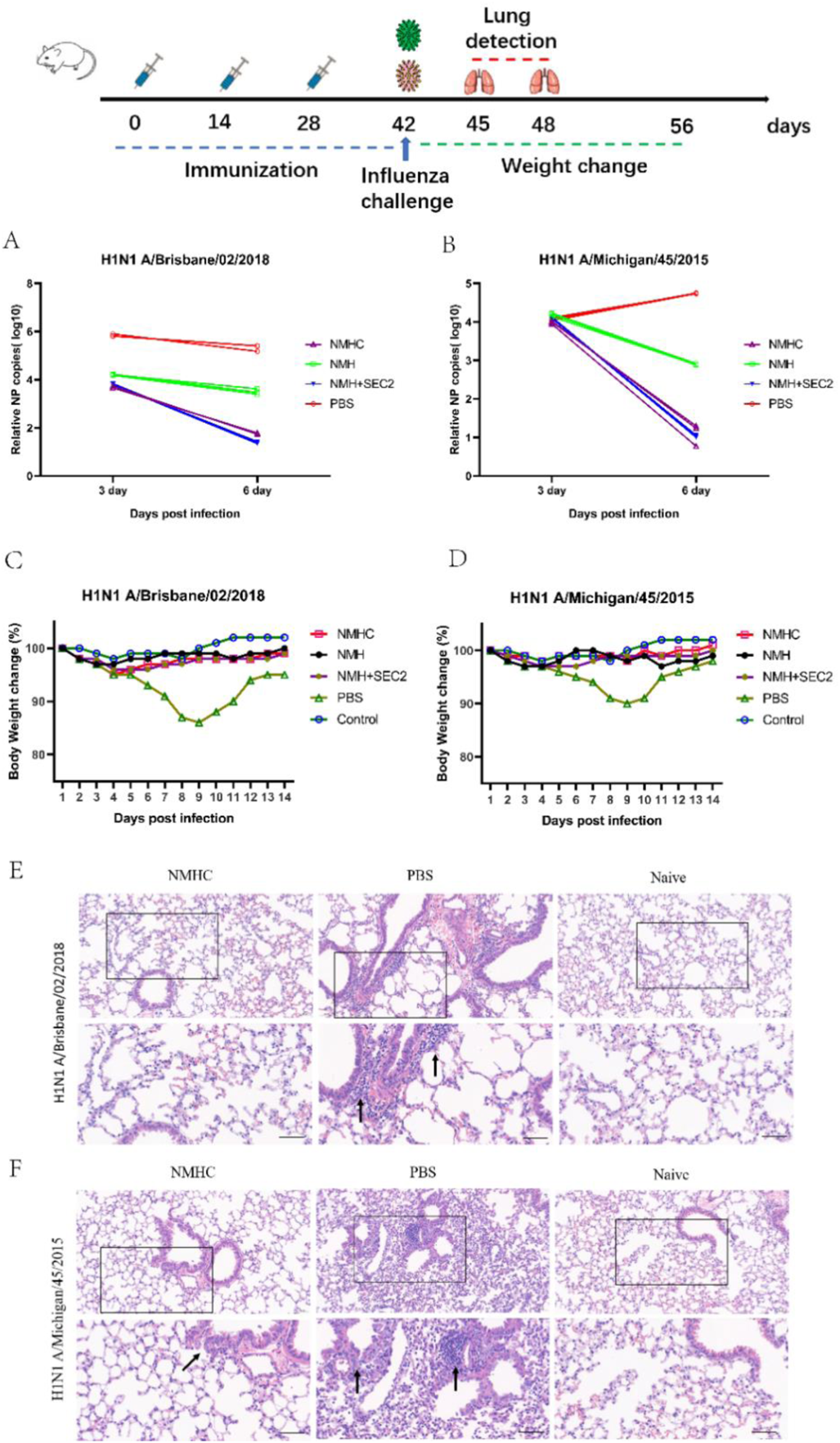
Challenge test in BALB/c mice. Mice (n=12 per group in two separate experiments) were infected intranasally with 10^3^ TCID_50_ of MI/45(H1) or Bris/02(H1) influenza viruses after 14 days of the third Immunization. A and B, viral copy number of MI/45(H1) and Bris/02(H1) in the lungs at day 3 and day 6 post-infection was determined using qPCR. C and D, weight changes (%) were monitored for 14 days post challenge. E and F, histopathology in pulmonary tissue (Scale bar = 50 μm). Naïve mice who were untreated and unchallenged were used as the control.

We examined the lymphocyte infiltration in the lungs of mice by immunohistochemistry method to evaluate the inflammatory damage caused by influenza virus infection. As expected, the leukocyte common antigen CD45 was widely detected in the lung sections of PBS treated mice but not in naïve mice without infection (*p* < 0.05 for Bris/02(H1) and *p* < 0.01 for MI/45(H1), Fig. 11). Immunized with NMHC before influenza virus infection significantly decreased the intensities of lung-infiltrated CD45 leukocytes compared to PBS treatment (Fig. 11B and C), which indicated a relieved inflammation damage in lung.

**Figure 11.**
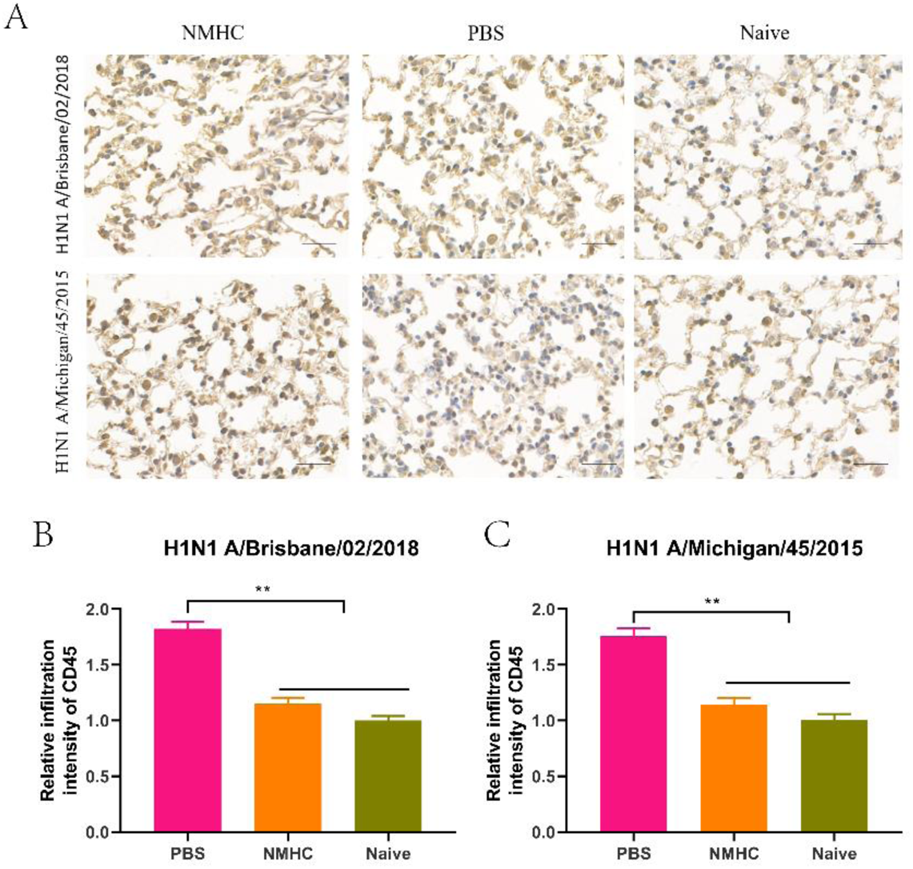
The profile analysis of lung-infiltrating leukocytes in lung frozen sections of infected mice treated with NMHC. A, lung-infiltrating leukocytes were analyzed by IHC of CD45. B and C, the quantification of the relative infiltration intensity of biomarkers by normalizing the positive signals of the treated groups to those of the naïve group. Error bars denote mean ± SD. **, *p* < 0.01. Scale bar = 50 μm.

Furthermore, immunofluorescence analysis of lungs sections with fluorescence Abs against the influenza nucleoprotein, macrophage common antigen (F4/80) and CD8 were performed to evaluate the distribution of influenza viruses, lung infiltrating macrophages, and lung infiltrating CD8^+^ CTL (Fig. 12A). As shown in Fig. 12B and C, at day 6 after infection, strong pink fluorescence signals of influenza nucleoprotein were detected in lungs of PBS treated mice, which indicated extensive distributions of influenza viruses. While the relative fluorescence intensities of influenza nucleoprotein in lungs of NMHC vaccinated mice were 4.5-fold (Bris/02(H1)) and 2.5-fold (MI/45(H1)) weaker than those of PBS treated mice. In both of the two infection models, NMHC immunization induced significantly enhanced distribution of CD8^+^ CTL in lungs compared to PBS control groups (*p* < 0.01), which were consistent with the results from CTL assay. After influenza virus infection, high levels of CD8^+^ CTL in lung lead to an effective clearance of virus infected cells, which contributed to the low virus loads as showed in nucleoprotein fluorescence signals. On the contrary, the intensities of macrophagocytes were significantly higher in PBS treated mice than those in naïve and NMHC immunized mice (*p* < 0.05), which might be attributable to heightened inflammation and delayed viral clearance in PBS treated mice(Valkenburg et al., 2014). This result verified that immunization with NMHC provided robust protection against heterologous virus challenge in vivo.

**Figure 12.**
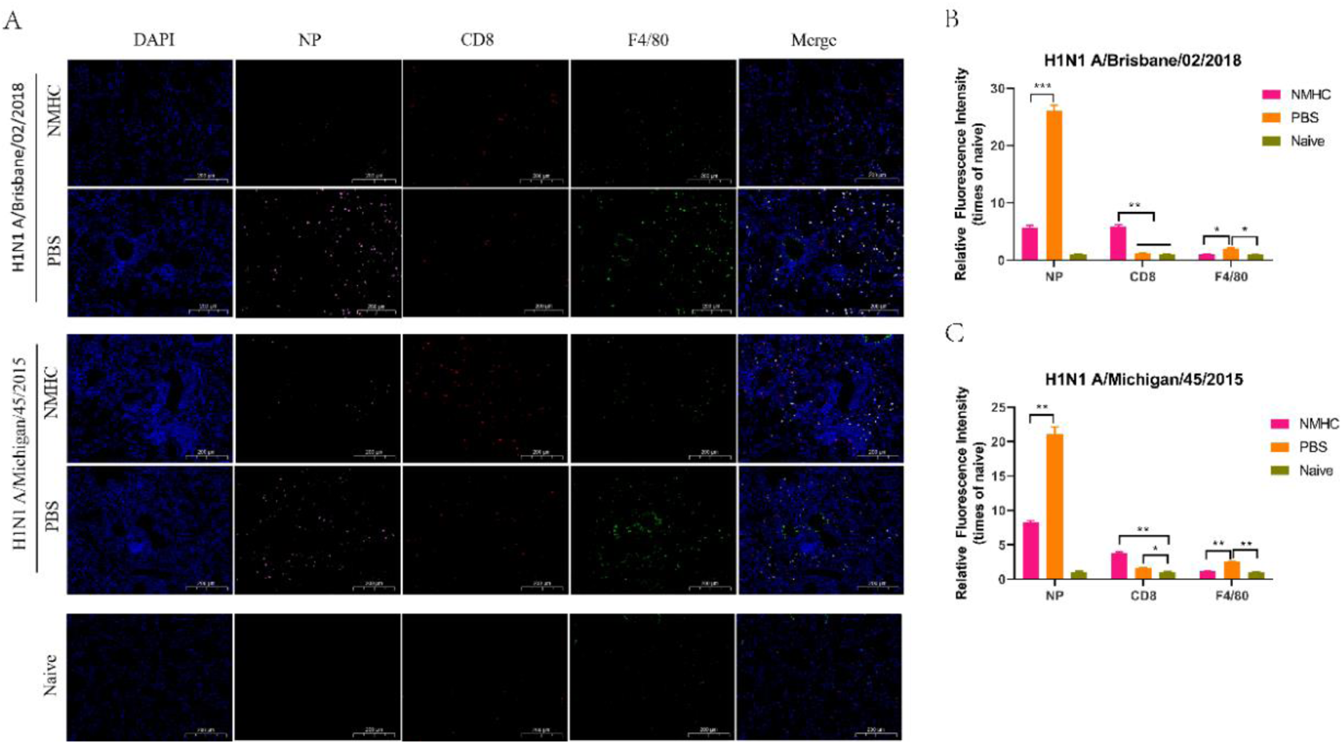
Immunofluorescence analysis of lungs frozen sections at day 6 after infection. A, distribution of influenza viruses (pink for NP), lung infiltrating CD8^+^ T cells (red for CD8) and lung infiltrating Macrophages (green for F4/80). B and C, the relative quantification of fluorescence intensity of NP, CD8 and F4/80. The relative fluorescence intensities were quantified by normalizing the fluorescent signals of the treated groups to those of the naïve group. Error bars denote mean ± SD. Scale bar = 200 μm, * *p* < 0.05, ** *p* < 0.01, *** *p* < 0.001.

## Discussion

In this study, we constructed a recombination protein NMHC as a novel universal influenza vaccine. NMHC consist of two domains, NMH domain as universal immunogen and SEC2 domain as adjuvant-like molecular chaperone, fused by a flexible peptide linker. Both of the two domains do not require post-translational modification, so NMHC can be conveniently and cheaply produced by *E. coli* expression system. Here, we studied humoral immunity responses mediated by NMHC involved in the anti-influenza im mune defense together with the contribution of cellular immunity. First and foremost, we demonstrated the protective mechanisms to influenza infection mediated by influenza specific humoral immunity responses. In vivo study, our data showed that NMHC could promote B cells to get recruited to the immune reaction and differentiate into NMH, H1N1, and H3N2 specific ASCs or memory cells in humoral immunity responses. Thereby, NMHC could obviously induce IgG2a and IgM but not IgE secretion with the number of immunizations increasing, and the antibody concentration reached peaks after the third immunization. During Th1-type immune response, IgG2a mediates clearance of virus and protection against influenza infection(Valkenburg et al., 2014), and IgM was regard as vanguard of initial humoral immunity, while IgE was associates with type I hypersensitivity. As expected, antiserum from NMHC immunized mice could specifically neutralize H1N1 and H3N2 so that the viruses could not replicate in MDCK cells as showed in Fig.4. In this experiment, virus suppression by antiserum were only detected by MN-HI assay but not by HI assay. The possible reason is that stem-specific antibodies did not bind to the head region of hemagglutinin, the receptor binding site (RBS) of influenza virus to target cell, and then did not prevent red-blood-cell aggregate induced by virus(He et al., 2015, van der Lubbe et al., 2018). Even if, influenza virus attached to the receptor and infected into cell by receptor-mediated endocytosis, and the stem-specific antibodies could bind to the non-receptor binding region of HA and prevent influenza virus from fusing into the endosomal membrane through interfering conformational change of HA induced by the low-pH (Fig. 13)(Imai et al., 1998). This procedure prevented the release of the ribonucleoprotein complex into the target cell and lead to the inhibition of viral replication, so the small number of viruses could not induce red-blood-cell aggregate which could be directly detected by HI assay. Next, in order to determine whether the antibody induced by NMHC had broad heterosubtypic binding or neutralization activity against diverse influenza A strains, we detected the binding ability of anti-NMHC sera to different HAs from group 1 (H1, H2, H5, H9) and group 2 (H3, H7) by ELISA. As expected, antiserum from NMHC immunized mice could broadly bind to these HA fragments with high antibody titers. Although only six kinds of HA were available in the materials of this experiment, we believed that the anti-NMHC serum could bind to more other HA subtypes. Additionally, results of SPR assay showed that antibodies purified from anti-NMHC serum exhibited high binding affinities to HAs of H1, H2, and H5, and moderate affinity to H3. Since the H3 fragment used in this study was from split virion of H3N2 but not from commercial purified full-length fragment, we believe that this relatively low affinity of antibodies from anti-NMHC serum to H3 might due to the impurity of the H3 fragment. These results verified that the antibody induced by NMHC vaccination could bind and neutralize varied influenza viruses with high affinities and effectively prevented virus replication in target cells.

**Figure 13.**
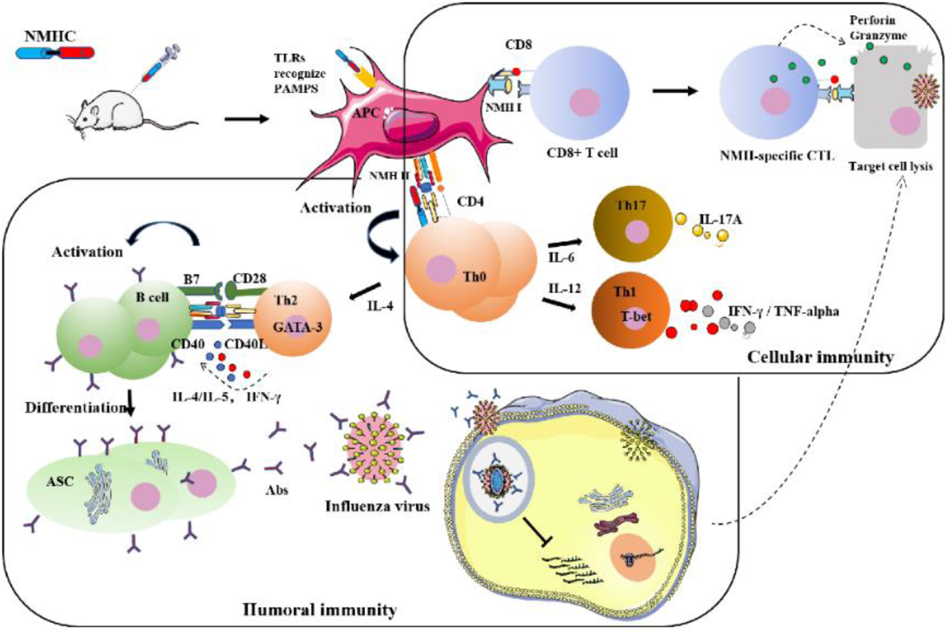
Detailed illustration of the anti-influenza mechanisms induced by NMHC.

Subsequently, in aspect of cellular immunity responses, we found that NMHC not only promoted T lymphocyte proliferation but also enhanced the functions of CD4^+^ and CD8^+^ T cells, respectively. CD4^+^ T cells can further differentiate into Th1 and Th2 cells which playing prominent role in immune responses to clear virus infection and to regulate immunoglobulin isotype switching(Snapper and Mond, 1993). Th1 can enhance IgG2a switching by secreting IFN-γ, and Th2 can enhance IgG1 switching by secreting IL-4. And IgG2a has been reported to be more effective than IgG1 in the resistance of virus infection(Yendo et al., 2016). CD8^+^ T cells can enhance antiviral responses by inducing the destruction of virus-infected cells though perforin and granzyme released by CTL. Our results from ELISPOT and qPCR assays indicated that immunization with NMHC significantly enhanced both Th1 and Th2 immune cells responses with increased IFN-γ and IL-4-producing cell differentiation and cytokines expression. Likewise, the antigen-specific CTL response was also elevated significantly, as showed in vivo CTL assay, the NMHC-immunized mice not only effectively killed target cells “infected” with NMH but not control cells “infected” with BSA, but also promoted the transcription of immune effector molecule of perforin and granzyme.

As expected, NMHC effectively induced the proliferation of murine splenocyte, which indicated that the recombinant protein NMHC maintained the superantigen activities of SEC2. Furthermore, NMHC significantly promoted the maturation of BMDCs with increasing expression of co-stimulatory molecules CD80, CD86 and MHC II in vitro, and we believe that this ability of NMHC is attributable to the combined effect of the NMH domain and SEC2 domain in the fusion protein, because CD80 and CD86 expressions in BMDCs induced by NMHC were significantly higher than those induced by SEC2, NMH, and NMH+SEC2, respectively. Matured BMDCs promoted CD4^+^ T cells to differentiate into immune response T cells subtype of Th1, Th2, and Th17 as represented by IFN-γ, IL-4, and IL-17A production. Compared with NMH and NMH+SEC2, this study implied that NMHC had an obviously adjuvant-like activity offered by the fused SEC2 domain, which could regulate the adaptive immune responses and augment the immunogenicity of NMH domain. The structural modeling of NMHC in silico revealed that the fused SEC2 was independent from the NMH region which benefited immune cells in recognizing components of antigenic determinants and inducing specific immune responses. Limited by laboratory biosafety, we only used two attenuated strains of H1N1 separated from different times and places, but not other virulent pathogenic strains, in mice challenge experiment. The result showed that immunization with NMHC could effectively protect mice against two virus strains infection according to the decreased number of viruses and relieved inflammation infiltration in the lungs. Although immunization with NMH+SEC2 achieved similar results to NMHC in reducing the number of viruses in lung, considering the comprehensive effects of humoral immunity and cellular immunity, as well as the convenience of future application, we believe that NMHC is more preferable than NMH+SEC2.

In summary (Fig. 13), recombinant vaccine NMHC could trigger both B cells and T cells immune responses against influenza virus infection. The SEC2 domain of NMHC could promote DCs maturation though TLR-NF-κB signaling pathways(Yao et al., 2018b), and matured DCs could effectively present the antigens of NMH domain to CD4^+^ T cell though MHC II molecule and CD8^+^ T cell though MHC I molecule. This process could lead to the activation, proliferation and differentiation of antigen-specific CD4^+^ T cells and CD8^+^ T cells. On the one hand, activated antigen-specific CD4^+^ T cells could further differentiate into IL-17A producing Th17 cells, IFN-γ producing Th1 cells and IL-4 producing Th2 cells. And Th1 and Th2 cells could help B cell to secret HA2 specific-antibodies which bind to HA2 of influenza virus and block the fusion of virus and endosomal membranes of target cells, so that prevent the release of ribonucleoprotein complex into the cytoplasm of target cells. On the other hand, with the aiding of Th1 cells, the activated antigen-specific CD8^+^ T cells could differentiate into CTLs effector cells which specifically kill virus-infected cells by exocytosis of cytolytic granules such as perforin and granzymes. Our findings imply an innovative potential clinical strategy for influenza immunization. We propose the recombinant protein NMHC as a potential candidate of universal broad-spectrum vaccine for various influenza virus prevention and therapy.

## Materials and Methods

### Animals

Female BALB/c mice (4–6weeks old) were purchased from Beijing Vital River Laboratory Animal Technology Co. Ltd (Beijing, China) and maintained under specific pathogen-free conditions. Feed and water were supplied ad libitum. All animal procedures were performed in accordance with Institutional Animal Care and Use Committee (IACUC) guidelines and have been approved by the IACUC of University of Chinese Academy of Sciences.

### Reagents

The expression vector pET-28a-sec2 containing full-length cDNA of SEC2 was constructed previously and preserved in our laboratory(Fu et al., 2017). Primers were synthesized by Sangon Biotech (Shanghai, China). RPMI 1640 and fetal bovine serum (FBS) were purchased from Thermo Fisher Scientific (Waltham, USA). IFN-γ and IL-4 enzyme-linked immune spot assay (ELISPOT) kits were purchased from eBioscience (Waltham, USA). Cell Titer 96 Aqueous One Solution Cell Proliferation assay (MTS) was purchased from Promega (Madison, USA). Goat anti-mouse IgG, IgG1, IgG2a peroxidase conjugate and all other reagents related to enzyme-linked immunosorbent assay (ELISA) were purchased from Bethyl Laboratories, Inc. (Montgomery, USA). CM5 chips for surface plasmon resonance (SPR) assay and Cytometric Bead Array (CBA) Mouse Immunoglobulin Isotyping Kit were purchased from GE HealthCare (Boston, USA). Ninety-six-well MultiScreen HTS plates were purchased from Millipore (Billerica, USA). RNAiso kit for total RNA extraction, PrimeScript RT Master Mix (Perfect Real Time) kit for RNA reverse transcription, and SYBR Premix Ex Taq Kit for real-time PCR assay were purchased from Takara (Dalian, China). Monoclonal CD3-FITC/APC, CD11c-APC, CD80-PE, CD86-FITC, MHC II-FITC, IL-4-FITC, IL-10-FITC, IL-17-PE, and IFN-γ-APC Antibodies (Abs) were purchased from BioLegend (San Diego, CA). CFSE Cell Tracking Kit was purchased from Beyotime Biotechnology (Haimen, Jiangsu, China) BD™. Beaver-Beads Protein A/G antibody Purification Kit was purchased from Beaver (Jiangsu, China). Commercial full-length HAs of H2N2 A/Canada/720/2005, H5N1 A/Hubei/1/2010, H7N9 A/Shanghai/2/2013 and H9N2 A/Hong Kong/35820/2009 were purchased from Sino Biological (Beijing, China). Viruses of H1N1 A/Michigan/45/2015, H1N1 A/Brisbane/02/2018 and H3N2 A/Hong Kong/ 4801/2014-like virus and split virion of H1N1 A/Michigan/45/2015 and H3N2 A/Hong Kong/ 4801/2014 were generously supplied by Chengda Biotechnology Co. Ltd (Shenyang, China).

### Construction and production of recombinant NMH and NMHC proteins

The recombinant protein NMH consists of two fragments of NP (335-350aa, 380-393aa) (Ben-Yedidia et al., 1999), two copies of M2e(Feng et al., 2006), and three tandem fragments of HA2 (76–130aa from H1N1 A/California/04/2009, H3N2 A/Hong Kong/1/1968, and H7N7 A/Netherlands/219/2003)(Wang et al., 2010). NMHC consists of NMH at the amino terminal followed by SEC2 sequence(Xu et al., 2008) at the carboxyl terminal. A flexible linker peptide sequence (GSAGSAG) was designed between each component to avoid any stereo-hindrance effect. Then, the encoding DNA fragment of NMH was optimized according to the codon preference of *E. coli* and synthesized by Sangon Biotech, while the DNA fragment of NMHC was constructed via overlap polymerase chain reaction (Overlap PCR). The encoding DNA fragment were ligated into the expression vectors pET-28a (+) plasmid and transformed into *E. coli* BL21(DE3), and the positive clones were verified by DNA sequencing. Vector-containing *E. coli* strains were cultured in LB (Luria-Bertani) medium, and protein expression was induced with 0.5 mM isopropyl β-D-1-thiogalactopyranoside (IPTG) at 30 ℃ for 8 h. Cells were harvested and disrupted by sonication on an ice bath, then supernatants and precipitates were isolated by centrifugation at 21,000 g for 15 min. SDS-PAGE electrophoresis results show that the expressed NMH and NMHC protein were both inclusion bodies. The method of protein purification and refolding was modified from previously reported (Lu et al., 2014, Song et al., 2020). The insoluble inclusion bodies were resuspended with washing buffer (2 M Urea, 50 mM Tris-HCl, 100 mM NaCl, 1 mM EDTA, pH 8.0), followed by centrifugation at 21,000 g for 15 min. This washing step was repeated twice before the washed inclusion bodies were solubilized in a denaturation buffer (8 M Urea, 100 mM NaH_2_PO_4_, 10 mM Tris-HCl, 20 mM imidazole, 1 mM DTT, pH 8.0) by votex-shaking at room temperature for 30 min. The supernatant was collected by centrifugation at 21,000 g for 15 min. The protein was purified by the Ni-saturated chelating sepharose affinity chromatography with the AKTA Fast Protein Liquid Chromatography System, eluting with a denaturation elution buffer (8 M Urea, 100 mM NaH_2_PO_4_, 10 mM Tris-HCl, 250 mM imidazole, 1 mM DTT, pH 8.0). To refold the elution fractions, dialysis refolding method was used. The refolding buffer was phosphate buffer saline (PBS) adding 0.5 M arginine, 0.2 mM oxidized glutathione (GSSG), 1 mM reduced glutathione (GSH) and 4 mM EDTA supplement with 6 M, 3 M, 1.5 M, 0.75 M, 0 M urea, respectively. The purified protein was refolded with step-wise dialysis proceeding from 6 M to 0 M urea following exchanging the refolding buffer three times with PBS.

### Recombinant protein-mediated splenocytes proliferation assay

Murine splenocytes were obtained from healthy BALB/c mice under aseptic condition as described in our previous report and maintained in RPMI 1640 medium containing 10% FBS. The freshly isolated murine splenocytes were seeded in 96-well flat-bottomed plates at 1 × 10^6^ cells/well and stimulated for 72 h with 3.5 μM NMHC, 3.5 μM NMH, 3.5 μM SEC2, and 3.5 μM NMH plus 3.5 μM SEC2, respectively, using PBS as negative controls. Cell proliferation was determined by MTS assay, and the proliferation index (PI) was calculated as described previously (Zhang et al., 2016).

### Recombinant protein-mediated BMDCs maturation and CD4^+^ T cell differentiation assay

Murine BMDCs were prepared as previously described(Yao et al., 2018b). Briefly, mice were sacrificed by intraperitoneal (i.p.) administration of tribromoethanol (400 mg/kg body weight), and the femur and tibia of the hind legs were dissected, then bone marrow cavities were flushed with 10 mL cold sterile PBS. After lysing red blood cells, the bone marrow cells were cultured and differentiated into BMDCs in RMPI-1640 with 10% FBS, 20 ng/mL rmGM-CSF, 10 ng/mL rmIL-4, 100 μg/mL streptomycin and 100 U/mL penicillin. Six days later, the purity of CD11c^+^ cells were > 90% as determined by flow cytometry. Then, BMDCs were simulated for 24 h with 3.5 μM NMH, 3.5 μM NMHC, 3.5 μM NMH plus 3.5 μM SEC2, and 3.5 μM SEC2, respectively, using PBS as negative control. The maturation of BMDCs were evaluated through detecting the cell surface markers including CD80, CD86, and MHC II with fluorescent labeled antibodies, respectively, followed by analyzing in a FACScalibur flow cytometer.

CD4^+^ T cells were sorted from freshly isolated murine splenocytes by immunomagnetic beads. For Th cells differentiation assay, the matured BMDCs were co-cultured with CD4^+^ T cells at a ratio of 10^4^:10^5^ for 96 h. Cells were fixed and permeabilized with Transcription Factor Buffer Set according to the manufacturer’s instructions. Then cells were intracellular stained with fluorescent labeled antibodies against IL-4 (for Th1 subtype), IFN-γ (for Th2 subtype), IL-17A (for Th17 subtype), and IL-10 (for Th10 subtype), respectively. The percentages of helper T cell subtypes were determined by flow cytometric.

### Animal Immunization

Female wild-type BALB/c mice were respectively immunized with an equimolar of NMHC (200 μg), NMH (100 μg), SEC2 (100 μg), NMH (100 μg) plus SEC2 (100 μg) and PBS (control) by i.p. (intraperitoneal) administration at day 0 (prime), 14 (boost) and 28 (boost). Serum samples were collected on the day 14, day 28, day 42, and day100 and kept at −80°C until use. Splenocytes were harvested at day 14 after the last immunization for ELISPOT assays and flow cytometric analysis.

### Mouse Immunoglobulin Isotyping detected by Cytometric Bead Assay

Isotype profiles of mouse immunoglobulins including IgA, IgE, IgG1, IgG2a, IgG2b, IgG3, and IgM were detected by Cytometric Bead Array (CBA) according to the manufacture’s instruction. Shortly, for serum sample diluted 4000 times in PBS, 50 μL Mouse Ig Capture Bead Array was mixed with 50 μL standards or sera samples diluted in master buffer (1:10,000) in tube before incubation for 15 minutes at room temperature. Then, the beads were washed once and added with 50 μL PE/FITC detector antibody in master buffer. After incubation for 15 minutes at room temperature in the dark, the beads were washed once and detected by flow cytometer. The mean fluorescence intensity (MFI) of PE represented the antibody expression levels.

### Microneutralization-Hemagglutinin inhibition assay (MN-HI)

Two strains of influenza virus were used for MN-HI assay to detect biological activity of serum antibody induced by recombinant protein(Fu et al., 2016). After treatment with receptor destroying enzyme, serum samples were serially double diluted in 96-well plates and mixed with H1N1 A/Michigan/45/2015 (MI/45(H1)) and H3N2 A/Hong Kong/ 4801/2014-like (HK/4801(H3)) at a final concentration of 100 TCID_50_ (Median Tissue Culture Infectious Doses) virus per well, respectively. After incubation for 1 h at 37°C, 1.5×10^4^ MDCK cells supplemented with 2 μg/mL trypsin and 0.5% Bovine Serum Albumin (BSA) in DMEM media were added to each well. After infection and amplification for 72 h at 37°C. Then viruses were collected and incubated with Turkey Red Blood cells in 96-well plate at room temperature for 30 min. The hemagglutination status of each well were visually determined. The titers of each serum sample were defined as the reciprocal of the highest dilution where no HI was observed.

### Measurement of virus-specific IgG by ELISA

The 96-well microtiter plates were coated with 100 μL HAs of H2N2 A/Canada/720/2005, H5N1 A/Hubei/1/2010, H7N9 A/Shanghai/2/2013, H9N2 A/Hong Kong/35820/2009 and split viruses of MI/45(H1), HK/4801(H3) at 2 mg/mL for overnight. Plates were then washed three times with washing buffer (PBS with 0.05% Tween-20, pH 7.4) and blocked with 200 μL blocking buffer (PBS with 0.05% Tween-20 and 1% BSA, pH 7.4) per well for 1 h at room temperature. After washing three times, 100 μL pre-serially diluted serum were added to each well and incubated at room temperature for 2 h. Plates were again washed five times before adding 100 μL secondary antibody (Horse radish peroxidase-labelled anti-mouse IgG, IgG1, IgG2a, 1:10,000 diluted in blocking solution) and further incubated for 1 h at room temperature. Then, the plates were washed four times and added with 100 μL TMB substrate per well for 10 min. Enzymatic color development was stopped with 100 μL of 2 M hydrochloric acid per well, and the plates were read at an absorbance of 450 nm. Titer was defined as the highest dilution of serum antibodies at which the mean OD_450_ value of the experiment group was no less than 2.1 times of the control. The experiment was repeated three times.

### Antibody Purification and Pull-Down Assay

Serum antibodies were purified using the Beaver-Beads Protein A/G antibody Purification Kit according to the protocols provided by the manufacturer. The ability of purified Abs to form a stable complex with NMH was further confirmed in a pull-down assay(Mallajosyula et al., 2014). NMH and Abs were mixed together at 2:1 molar ratio and incubated for 2 h at 4°C. The equilibrated Protein A/G beads were added to the mixture and incubated for 2 h to bind and pull down NMH, while the unbound supernatants were separated. The antibody bound to the beads were eluted with antibody elution buffer and then neutralized with antibody neutralization buffer. The unbound and eluted fractions were subsequently analyzed by SDS-PAGE.

### Binding Affinity Studies Using SPR

Binding affinities of the serum antibody with HAs of H2N2, H5N1, and split virion of H1N1, H3N2 as described above were determined by SPR experiments performed with Biacore T200(Fu et al., 2016). The purified Abs in PBSP buffer (PBS with 0.05% P20, pH 7.4) at 1.25 nM were captured onto CM5 chip though goat anti-mouse IgG (H+L) which immobilized (five hundred to seven hundred response units (RU)) by standard amine coupling to the surface of the biosensors. The goat anti-mouse IgG (H+L) sensor channel served as a negative control for each binding interaction. Multiple concentrations of HAs were passed over each channel in a running buffer of PBS (pH 7.4) with 0.05% P20 surfactant. Both binding and dissociation events were measured at a flow rate of 30 μL/min. The sensor surface was regenerated after every binding event by repeated washing with Glycine pH 2.0. Each binding curve was analyzed after correcting for nonspecific binding by subtraction of signal obtained from the negative control flow channel. The KD values were calculated using a steady affinity state model by the BIAcore T200 evaluation software (Version 3.1)(Zhang et al., 2013).

### ELISPOT assays

Splenocytes were freshly isolated from immunized mice of each group. Influenza-specific IgG antibody secreting cells (ASCs) were enumerated with ELISPOT. MultiScreen HTS 96-well plates were coated with purified NMH at a concentration of 5 μg/mL in PBS to detect antigens specific ASCs, or coated with split viruses of H1N1 and H3N2 at a concentration of 5 μg/mL in PBS to detect influenza virus-specific ASCs. Wells coated with PBS were served as negative controls. Plates were incubated overnight at 4°C and blocked with complete medium for 2 h at 37°C. Then splenocytes were added into triplicate wells (500,000 cells/well) and incubated at 37°C for 20 h. Plates were vigorously washed to remove cells, and then incubated with HRP-conjugated anti-mouse IgG overnight at 4°C. The spots representing the IgG ASCs in each well were counted by inspection. Nonspecific spots detected in the negative control (PBS) wells were subtracted from the counts of each total ASCs. Moreover, ELISPOT kits were used to measure the IFN-γ or IL-4-producing-splenocytes onsite according to the manufacturer’s protocol.

### Real-time Quantitative polymerase chain reaction (qPCR)

To detect inflammatory cytokines, transcription factor expression in splenocytes and the virus titers in lungs of infected mice, total RNA were extracted from splenocytes or lung tissues using the RNA-extracting reagent RNAiso Plus, and 1 μg of total RNA were reverse transcribed to cDNA using a PrimeScript RT Master Kit according to the manufacturer’s instructions. Resulting cDNA was used for quantitative real-time PCR (qPCR) analysis with a SYBR Green PCR kit (Roche). β-Actin was used as reference gene. All primers were listed in Table 1. Relative transcription levels were determined using the 2^−ΔΔCt^ analysis method(Fu et al., 2020).

### Cytotoxic lysis assay

In vivo cytotoxic lysis assay was performed as previously reported(Xie et al., 2014). Splenocytes from naïve BALB/c mice were divided into 2 parts. One part was pulsed with 10^−6^ M NMH peptides and labeled with 10 μΜ of CFSE (defined as CFSE^high^ target cells). The other part was pulsed with 10^−6^ M BSA and labeled with 1 μM of CFSE (defined as CFSE^low^ cells) as a non-target control. Cells from the 2 parts were mixed in a 1:1 ratio and injected into immunized recipient mice at 2 × 10^7^ total cells per mouse via the tail vein on day 14 after the third immunization. Twelve hours later, splenocytes were isolated from the recipients and differential CFSE fluorescent intensities were measured with a flow cytometry. Specific lysis was calculated using the following formula: Percentage of specific lysis= (1-[ratio unprimed/ratio primed] × 100), where ratio = percentage CFSE^low^/percentage CFSE^high^.

### Challenge experiments

The challenge experiments were performed as previously reported(Meng et al., 2013). BALB/c mice were immunized with 200 μg NMHC, 100 μg NMH alone, 100 μg NMH plus 100 μg SEC2, or PBS. Two weeks after the third immunization, mice were challenged with virus. Before virus infection, 12 mice of each group were anesthetized and inoculated intranasally with 10^3^ TCID_50_ of MI/45(H1) and H1N1 A/Brisbane/02/2018 (Bris/02(H1)) virus strains in a volume of 50 μL. To determine the viral load and the pathological damage in the infected lungs, 3 mice from each group were sacrificed on day 3 and day 6 post-infection. As previously reported(Meng et al., 2013, Marcos et al., 2017), the right lungs were used for qPCR to assess the virus titer, while the left lungs were fixed for the histopathological analysis. Survival and weight change of the remaining mice in each group were monitored daily for 14 days after the infection.

### Histological analysis

The mice were sacrificed on day 3 and 6 after infection with virus. The left lungs were collected and dissected for histological observation (n = 3 mice per group). After fixation in neutral-buffered fixative, the tissues were embedded in paraffin and stained with hematoxylin and eosin (H&E). The lungs were sliced into 6-μm thick frozen sections and the lung-infiltrating leukocytes profiles in the tissues were reflected by IHC analysis of CD45. The distribution of influenza viruses, lung-infiltrating T lymphocytes and lung-infiltrating macrophages were respectively reflected by Immunofluorescence analysis of NP, CD8, and F4/80 with Laser Scanning Confocal Microscope (LSCM).

### Statistical analysis

The data were analyzed by Student’s t-test, one-way analysis of variance (ANOVA) and followed by a suitable post hoc test using the SPSS 26.0 and GraphPad Prism Software (version 6.0c). Differences with *p* values < 0.05 considered to be statistically.

### Funding

This work was supported by Strategic Priority Research Program of the Chinese Academy of Sciences Grant (XDA12020225), Liaoning Revitalization Talents Program (XLYC1807226), Science and Technology Plan Projects of Shenyang City Grants (Z17-7-013), and Shenyang High-level Innovative Talents Program (RC190060).

### Source data

Figure 1—source data 1: Source file for the gel data used for the qualitative analyses of NMHC, SEC2 and NMH purified and renatured from BL21(DE3) lysate shown in Fig. 1. This folder contains the original files of the full raw unedited gel (named original gel), and the relevant bands clearly labelled gel (named labelled gel). The information can be found in figure 1 legend, as well as methods.

Figure 6—source data 2: Source file for affinities data shown in Fig. 6. This folder contains the overlays of binding kinetics of antibodies to HAs (HA of H1, H2, H3 and H5) at different concentrations were detected with Biacore (individual files are named “the binding kinetics of antibodies to H1.txt”, “the binding kinetics of antibodies to H2.txt”, “the binding kinetics of antibodies to H3.txt”, “the binding kinetics of antibodies to H5.txt”). The detection processes are contained in the methods section.

Figure supplement 1-source data 3: Source file for pull-down assay shown in Fig. S1. This folder contains the original files of the full raw unedited gel (individual files are named “original gel (Line1-6)”, “original gel (Line7-15)”, “original gel (Line16-20)”), and the relevant bands clearly labelled gel (individual files are named “labelled gel (Line1-6)”, “labelled gel (Line7-15)”, “labelled gel (Line16-20)”). The information can be found in figure supplement 1 legends, as well as methods.

## Supplementary Information

**Figure S1.**
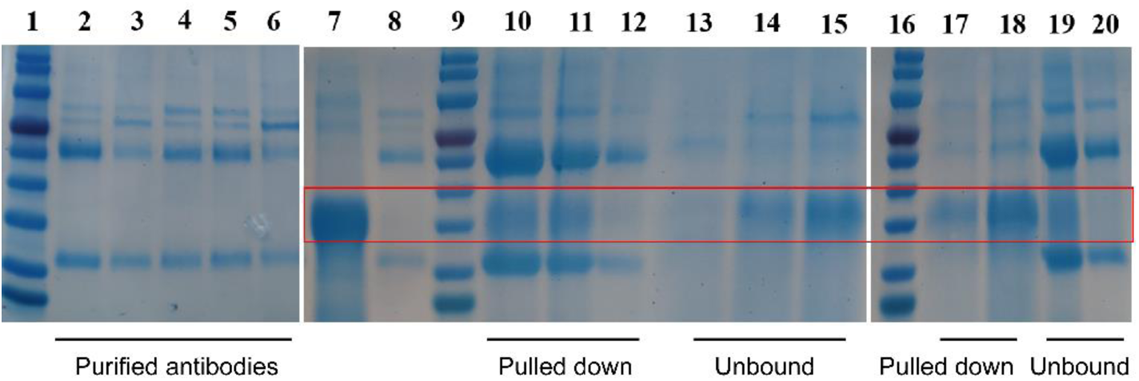
NMH forms a stable complex with the treated serum. NMH and treated serum were mixed together at 2:1 molar ratio and incubated for 2 h at 4 °C. Protein A/G beads specific for the IgG could pull down the antibody–antigen (Ab–Ag) complex. The Protein A/G-bound protein or not was eluted (elution fraction) with 100 mM glycine-HCl (pH 3). Lanes: 1,9 and 16 markers; lanes 2-6: pure serum of group treated NMHC, NMH, SEC2, NMH+SEC2 and PBS; lane 7: NMH; lanes 8: pure serum; lanes 10-12: elution fraction of NMHC, NMH and PBS; 13-15,17-18 lanes: unbound fraction; lanes 19 and 20: elution fraction of NMH+SEC2 and SEC2. All of the samples (with reducing agent) were analyzed on a denaturing SDS/PAGE. Source files of the gels used for the qualitative analyses were available in the Figure supplement 1-source data 3.

**Table S1.** Neutralizing antibody titers detected by standard HI

| Immunization |  | Serum dilution(1/X) |  |  |  |
| --- | --- | --- | --- | --- | --- |
|  |  | H1N1 | H3N2 | Victoria | Yamagata |
| Day14 | NMHC | < 1:8 | < 1:8 | < 1:8 | < 1:8 |
|  | NMH | < 1:8 | < 1:8 | < 1:8 | < 1:8 |
|  | SEC2 | < 1:8 | < 1:8 | < 1:8 | < 1:8 |
|  | NMH+SEC2 | < 1:8 | < 1:8 | < 1:8 | < 1:8 |
|  | PBS | < 1:8 | < 1:8 | < 1:8 | < 1:8 |
|  | QIV | 1:128 | 1:16 | 1:64 | 1:32 |
| Day28 | NMHC | < 1:8 | < 1:8 | < 1:8 | < 1:8 |
|  | NMH | < 1:8 | < 1:8 | < 1:8 | < 1:8 |
|  | SEC2 | < 1:8 | < 1:8 | < 1:8 | < 1:8 |
|  | NMH+SEC2 | < 1:8 | < 1:8 | < 1:8 | < 1:8 |
|  | PBS | < 1:8 | < 1:8 | < 1:8 | < 1:8 |
|  | QIV | 1:256 | 1:256 | 1:256 | 1:256 |
| Day42 | NMHC | < 1:8 | < 1:8 | < 1:8 | < 1:8 |
|  | NMH | < 1:8 | < 1:8 | < 1:8 | < 1:8 |
|  | SEC2 | < 1:8 | < 1:8 | < 1:8 | < 1:8 |
|  | NMH+SEC2 | < 1:8 | < 1:8 | < 1:8 | < 1:8 |
|  | PBS | < 1:8 | < 1:8 | < 1:8 | < 1:8 |
|  | QIV | 1:256 | 1:25 | 1:512 | 1:256 |

Quadrivalent influenza vaccine (QIV) was regarded as positive control.

**Table S2.** ELISA endpoint titers of HAs or split virion

| Vaccine | Mouse | Titer against |  |  |  |  |  |  |  |  |  |
| --- | --- | --- | --- | --- | --- | --- | --- | --- | --- | --- | --- |
|  |  | H1N1 |  |  | H3N2 |  |  | H2N2 | H5N1 | H7N9 | H9N2 |
|  |  | IgG | IgG1 | IgG2a | IgG | IgG1 | IgG2a | IgG | IgG | IgG | IgG |
| NMHC | No.1 | 3200 | 1600 | 400 | 3200 | 1600 | 400 | 8192 | 8192 | 8192 | 8192 |
|  | No.2 | 6400 | 3200 | 400 | 3200 | 1600 | 200 | 8192 | 8192 | 8192 | 8192 |
|  | No.3 | 6400 | 3200 | 800 | 3200 | 800 | 200 | 4096 | 8192 | 8192 | 8192 |
| NMH | No.1 | 1600 | 1600 | 100 | 400 | 400 | 200 | 8192 | 2048 | 2048 | 4096 |
|  | No.2 | 1600 | 1600 | 100 | 400 | 400 | 200 | 8192 | 4096 | 2048 | 4096 |
|  | No.3 | 1600 | 1600 | 100 | 400 | 400 | 200 | 4096 | 4096 | 2048 | 8192 |
| SEC2 | No.1 | <10 | <10 | <10 | <10 | <10 | <10 | <10 | <10 | <10 | <10 |
|  | No.2 | <10 | <10 | <10 | <10 | <10 | <10 | <10 | <10 | <10 | <10 |
|  | No.3 | <10 | <10 | <10 | <10 | <10 | <10 | <10 | <10 | <10 | <10 |
| NMH<br>+SEC2 | No.1 | 6400 | 3200 | 100 | 1600 | 1600 | 100 | 8192 | 8192 | 4096 | 8192 |
|  | No.2 | 6400 | 3200 | 100 | 3200 | 3200 | 100 | 8192 | 8192 | 4096 | 8192 |
|  | No.3 | 6400 | 6400 | 100 | 3200 | 3200 | 200 | 8192 | 4096 | 4096 | 8192 |
| PBS | No.1 | <10 | <10 | <10 | <10 | <10 | <10 | <10 | <10 | <10 | <10 |
|  | No.2 | <10 | <10 | <10 | <10 | <10 | <10 | <10 | <10 | <10 | <10 |
|  | No.3 | <10 | <10 | <10 | <10 | <10 | <10 | <10 | <10 | <10 | <10 |

